# Knock-out of Tpm4.2/actin filaments alters neuronal signaling, neurite outgrowth and behavioral phenotypes in mice

**DOI:** 10.1101/2025.04.10.648285

**Authors:** Sian Genoud, Chanchanok Chaichim, Rossana Rosa Porto, Tamara Tomanic, Holly Stefen, Esmeralda Paric, Soumalya Sarkar, Dasol Yoo, Wendi Gao, Edna C. Hardeman, Peter W. Gunning, Tim Karl, John Power, Thomas Fath

**Affiliations:** Faculty of Medicine, Health & Human Sciences, Macquarie Medical School, Dementia Research Centre, Macquarie University, Sydney, NSW, Australia; School of Medical Sciences, University of New South Wales, Sydney, NSW, Australia; School of Medicine, Western Sydney University, Campbelltown, NSW 2560, Australia

**Keywords:** Tyopomyosin, actin cytoskeleton, neurons, neuronal signaling

## Abstract

Tropomyosins (Tpm) are master regulators of actin dynamics through forming co-polymers with filamentous actin. Despite the well-understood function of muscle Tpms in the contractile apparatus of muscle cells, much less is known about the diverse physiological function of cytoplasmic Tpms in eukaryotic cells. Here, we investigated the role of the Tpm4.2 isoform in neuronal processes including signaling, neurite outgrowth and receptor recycling using primary neurons from Tpm4.2 knock-out mice. Live imaging of calcium and electro-physiology data demonstrated increased frequency, yet reduced strength of single neuron spikes. Calcium imaging further showed increase in neuronal networks. *In vitro* assays of Tpm4.2 knock-out neurons displayed impaired recycling of the AMPA neurotransmitter receptor subunit GluA1. Morphometric analysis of neurite growth showed increased dendritic complexity and altered dendritic spine morphology in Tpm4.2 knock-out primary neurons. Behavioral analysis of Tpm4.2 knock-out mice displayed heightened anxiety in Open field test whilst Elevated Plus maze displayed heightened anxiety only in females. A sex-dependent phenotype was also seen in the Social Preference test with impaired social memory and socialization in female Tpm4.2 knock-out mice, whilst male and female knock-out mice had impaired recognition memory in a Novel Object Recognition test. Our study depicts the multi-faceted role of the Tpm4.2 isoform and its co-polymer F-actin population in neurons, with potential implications for better understanding diseases of the nervous system which involve actin cytoskeleton dys-function.

## 1. Introduction

Tropomyosins (Tpm) are helical coiled-coil dimers that polymerize head-to-tail along the actin filament and are regarded as a master regulator of actin dynamics in mammalian cells as they regulate the access of other actin-binding proteins [1]. The co-polymer nature of Tpms with filamentous actin (F-actin) is suggested to determine specific F-actin functions through influencing F-actin stability and promoting or inhibiting the activity of other actin-binding proteins [2,3]. As Tpm isoforms are associated with molecularly distinct F-actin populations, Tpm isoforms can be used as a proxy to investigate functional properties of F-actin populations [4]. In eukaryotic cells, over 40 different Tpm isoforms are found, arising by alternative splicing from four different genes. The expression of these isoforms has been found to be spatially and temporally regulated. Products from Tpm1, 3 and 4 are expressed in neuronal cells and have distinct spatial and temporal distributions related to their distinct functions [5]. Little is known regarding the physiological function of Tpm4 (also previously known as deltaTm) in non-muscle cells, however Tpm4.2 is the only identified Tpm4 isoform in mice [6] and has been shown to enhance non-muscle myosin recruitment [7] and facilitate ER/Golgi trafficking. [8].

Previous studies have identified Tpm4 isoforms expressed in neuron-like cells early in development, before a reduction in expression levels during maturation [9]. During neuronal differentiation, there is a shift from enrichment in axonal growth cone [10], to enrichment in the post-synaptic density [11], suggesting a functional change from a potential role in neurite outgrowth to a proposed role in synaptic plasticity of mature neurons [12,11]. This is supported by evidence of increased neurites, filopodia, branch formation and enlarged growth cone area in differentiating B35 neuroepithelial cells, overexpressing Tpm4.2. [13]. Overexpression of Tpm4.2 is associated with an increase in phosphorylated (inactive) ADF(actin depolymerizing factor)/cofilin which act by severing actin filaments, therefore leading to increased actin stability and filament length [4,13]. As the post-synaptic compartment contains a complex of scaffolding proteins, receptors, actin-cytoskeleton proteins, adhesion and signaling molecules, understanding the physiological function of Tpm4.2 in this compartment could provide essential insight into neuronal growth and synapse formation and signaling. In this study, we aim to further elucidate the physiological function of Tpm4.2 in neuronal development, maintenance and signaling using primary neurons and behavioral testing in Tpm4.2 knock-out (Tpm4.2^-/-^) mice [6].

## 2. Results

### 2.1 Expression profile of Tpms in mouse brain

Immunoblotting was used to assess the brain region-specific expression of Tpm4.2 in the olfactory bulb, hippocampus, cerebellum, anterior cortex and posterior cortex of 7 months old wild-type (Tpm4.2^+/+^) mice. Tpm4.2 was similarly expressed in all brain regions with a trend towards higher expression levels found in the cerebellum. However, there was no significant difference in the relative abundance of Tpm4.2 between all brain regions (p > 0.05; **Figure 1**). Brain region-specific expression of Tpm4.2 was further assessed in the original C57Bl6 strain. This revealed significantly higher expression levels of Tpm4.2 in the cerebellum, compared to the hippocampus, anterior cortex and posterior cortex (**Figure S1**). Relative abundance normalized to total protein is outlined in **Table S1.**

**Figure 1.**
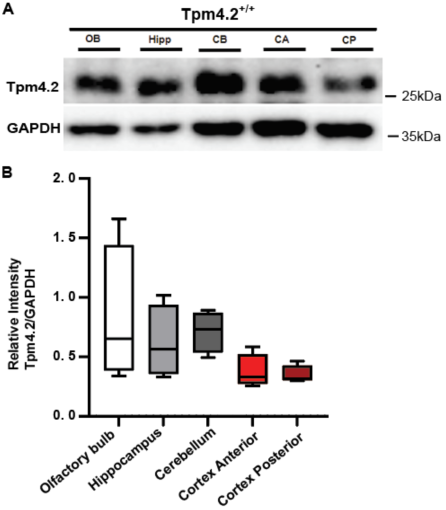
Tpm4.2 expression in the Tpm4.2^+/+^ mouse brain. **A** Immunoblot results of sub-dissected brains from Tpm4.2^+/+^ mice probed for Tpm4.2 (28kDa) and normalized to Glyceraldehyde 3-phosphate dehydrogenase (GAPDH; 36kDa). Olfactory bulb (OB), Hippocampus (Hipp), Cerebellum (CB), Anterior cortex (CA), Posterior cortex (CP). **B** Data are represented as min-max box plots with n = 4 per group. One-Way ANOVA followed by Bonferroni’s multiple comparisons detected no significant difference between brain regions (p > 0.05). ns = not significant.

**Figure S1.**
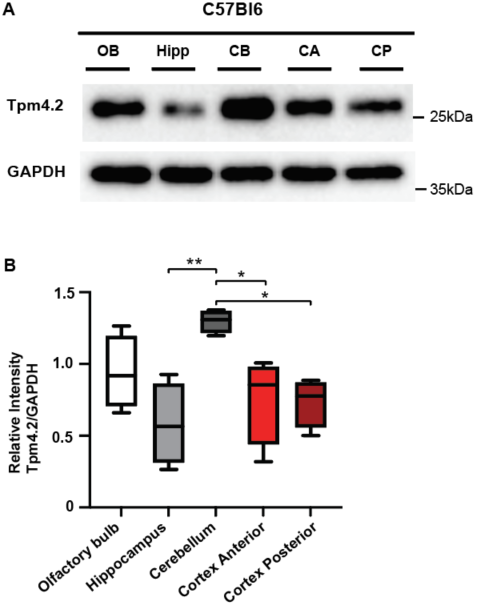
Assessing Tpm4.2 brain region-specific expression in C57Bl6 strain mice. **A** Immunoblots, probing for Tpm4.2 and GAPDH. Data are represented as min-max box plots, n = 4 per group, and statistical analysis, using One-way ANOVA, followed by Tukey’s multiple comparisons. Olfactory bulb (OB), Hippocampus (Hipp), Cerebellum (CB), Anterior cortex (CA), posterior cortex (CP). B There was significantly higher expression of Tpm4.2 in the cerebellum, compared to hippocampus, anterior and posterior cortex (p > 0.05).

**Table S1.**
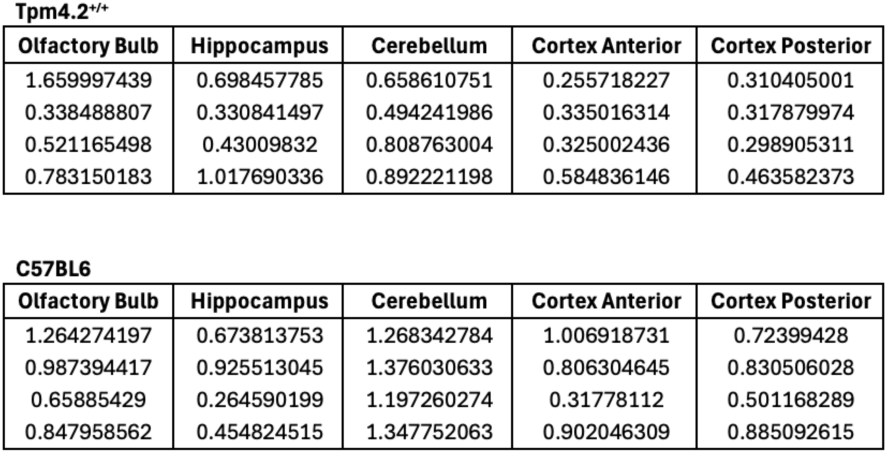
Relative abundance of Tpm4.2 in brain regions of Tpm4.2^+/+^ and C57Bl6 strain mice.

To determine if any other Tpm isoforms compensate for Tpm4.2 deletion, immunoblotting was performed on brain tissue from 7-months-old Tpm4.2^+/+^ and Tpm4.2^-/-^ mice. There were no significant differences in the relative expression of the Tpm3.1/2 isoforms in Tpm4.2^+/+^ (1.051 ± 0.082) and Tpm4.2^-/-^ (1.078 ± 0.056), in total *Tpm3* products between Tpm4.2^+/+^ (0.17 ± 0.020) and Tpm4.2^-/-^ (0.22 ± 0.018), or in Tpm1.10/12 isoforms between Tpm4.2^+/+^ (1.017 ± 0.054) and Tpm4.2^-/-^ (0.98 ± 0.036) mice (**Figure S2**).

**Figure S2.**
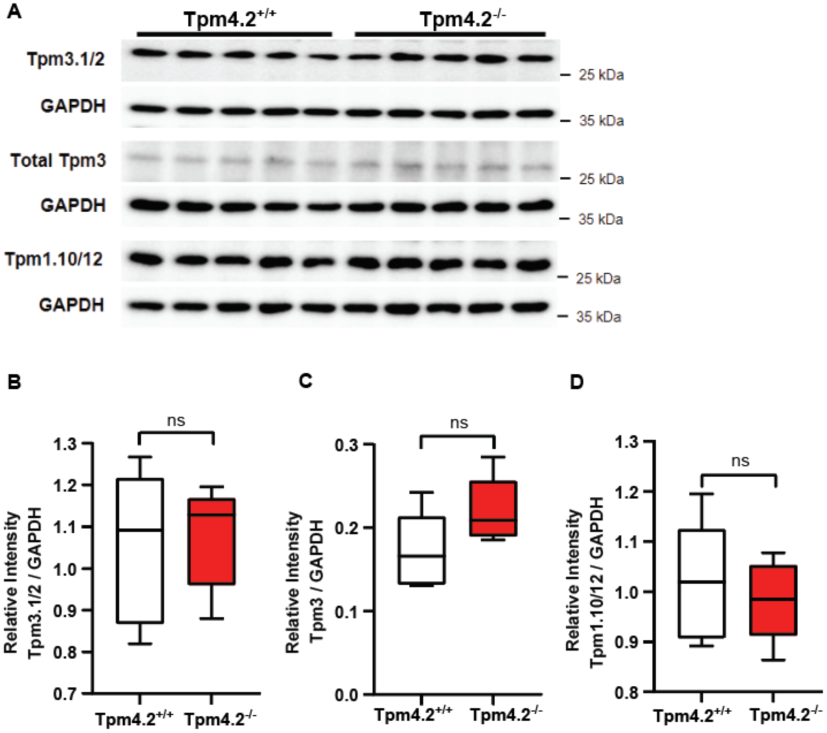
Assessing compensation of other Tpm isoforms in Tpm4.2^-/-^ mouse brains. **A** Immunoblots, probing for Tpm3.1/2, total Tpm3 isoforms and Tpm1.10/11 in both Tpm4.2^+/+^ and Tpm4.2^-/-^ mouse brains. **B-D** Data are represented as min-max box plots, n = 5 per group, and statistical analysis, using unpaired t-tests. There was no significant difference in the relative abundance of **B** Tpm3.1/2, **C** total Tpm3 isoforms or **D** Tpm1.10/12 in Tpm4.2^-/-^, compared with Tpm4.2^+/+^ brains (p > 0.05). ns = not significant.

### 2.2 GluA1 receptor recycling in Tpm4.2^-/-^ neurons

To assess whether Tpm4.2 plays a role in receptor recycling pathways, we performed a receptor internalization assay probing for surface and total (surface plus internalized) GluA1 receptors on dendrites of Tpm4.2^+/+^ and Tpm4.2^-/-^ neurons following stimulation with glycine, N-methyl-D-aspartate (NMDA) and bicuculline (**Figure 2A, B)**. As expected, stimulated Tpm4.2^+/+^ neurons showed a reduction in GluA1 receptors on the surface, compared with unstimulated neurons (**Figure 2C**). NMDA stimulated neurons had a 43.65% reduction in surface:total receptor ratio, when compared with control extracellular solution (ECS) incubated neurons (p = 0.042). Glycine stimulation reduced this ratio by 63.21% (p = 0.0092), and bicuculline-stimulated neurons had a 49.84% reduction in surface:total receptors, when compared with ECS (p = 0.011). Tpm4.2^-/-^ neurons had a much higher surface:total GluA1 receptor ratio when stimulated with NMDA (52.77% increase; p = 0.0035), glycine (66.45% increase; p = 0.0002) and bicuculline (44.79%; p = 0.012, **Figure 2C**).

**Figure 2.**
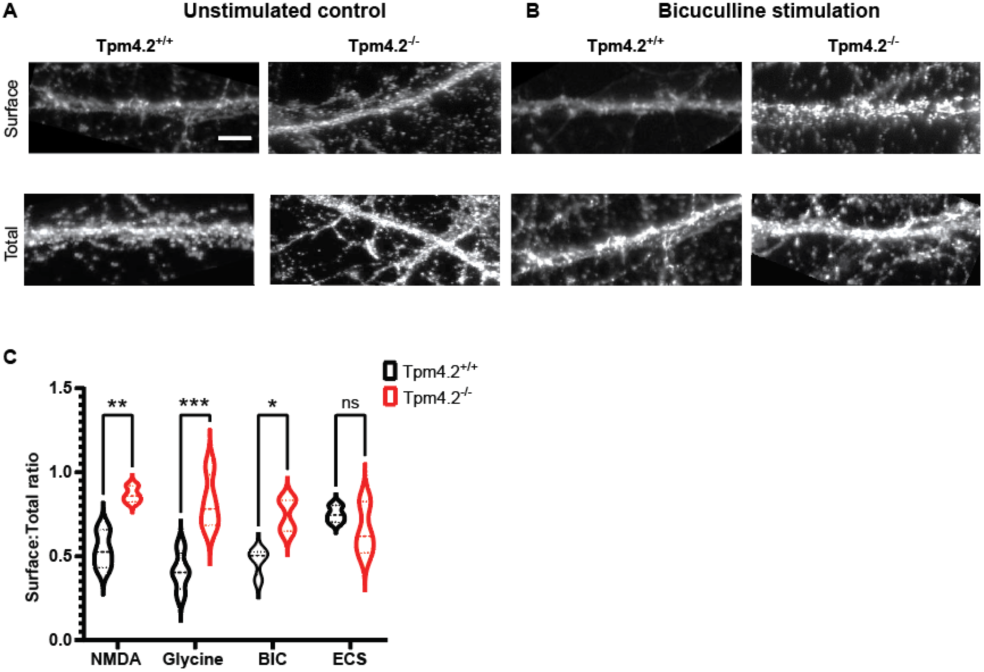
GluA1 receptor recycling in Tpm4.2^-/-^ neurons. **A** Representative immunocytochemistry images of surface and total GluA1 receptors from unstimulated 18 days in vitro primary hippocampal Tpm4.2^+/+^ and Tpm4.2^-/-^ neurons in extracellular solution (ECS), compared with **B** Surface and total GluA1 receptors on Bicuculline stimulated neurons **C** A significant increase in surface to total (surface plus internalized) ratio was observed in NMDA, glycine and bicuculline (BIC) stimulated neurons. GluA1 receptors of Tpm4.2^-/-^, compared with Tpm4.2^+/+^ neurons, reflective of a loss in GluA1 internalization. n= 10 dendrites per neuron, 10 neurons per condition x 4 biological replicates. Red boxes indicate area of dendrite measured. Data are represented as min-max violin plots with median and quartiles indicated by solid and dashed lines respectively. Scalebar = 5µm ns = not significant, *p < 0.05, **p < 0.005, ***p < 0.0005, ****p < 0.0001.

### 2.3 Neuronal activity in Tpm4.2^-/-^ versus Tpm4.2^+/+^ neurons

#### 2.3.1 Calcium Spike Analysis

Fluorescent calcium imaging was used to assess the role of Tpm4.2 in controlling neural network activity. Tpm4.2^+/+^ and Tpm4.2^-/-^ neuronal cultures were imaged at 20 days *in vitro* (DIV) (Tpm4.2^-/-^: n = 2094 neurons / 26 FOV / 4 preparations; Tpm4.2^+/+^: n = 775 neurons / 7 FOV / 3 preparations). Neuronal density did not differ between groups (p > 0.05, **Figure S3**).

**Figure S3.**
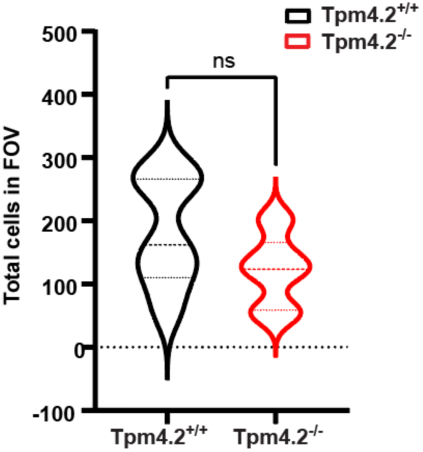
Cell count of cells per field of view (FOV), that were analyzed by calcium imaging. There was no significant difference in the cell density of Tpm4.2^+/+^ and Tpm4.2^-/-^ neurons in the fields of view that were analyzed for calcium spikes.

Neuronal activity was primarily characterized by synchronized calcium spikes (**Figure 3A(a), A(b)**. These synchronous events were on average 138% more frequent in the Tpm4.2^-/-^ (3.1 ± 0.3 Hz) than in Tpm4.2^+/+^ (1.3 ± 0.5 Hz) neuronal cultures (unpaired t-test, t(31) = 2.816, p = 0.008; **Figure 3B**). The amplitude of the synchronous events did not differ between Tpm4.2^-/-^ and Tpm4.2^+/+^ cultures (Mann-Whitney U = 76, p = 0.53; **Figure 3C**). Calcium spikes in individual neurons occurred 65% more frequently in the Tpm4.2^-/-^ (3.9 Hz) than in Tpm4.2^+/+^ (2.3 Hz) neurons (nested t-test, t(31) = 2.251, p = 0.03; **Figure 3D**). However, these individual spike peaks were ∼60% smaller in Tpm4.2^-/-^ (0.08 ΛιF/F) than Tpm4.2^+/+^ (0.20 ΛιF/F) neurons (nested t-test, t(31) = 2.845, p = 0.008; **Figure 3E**). No differences were observed in the coefficient of variation of either the amplitude (nested t-test, t(31) = 1.037, p = 0.31) or interevent interval (nested t-test, t(31) = 1.037, p = 0.31) of the calcium spikes.

**Figure 3.**
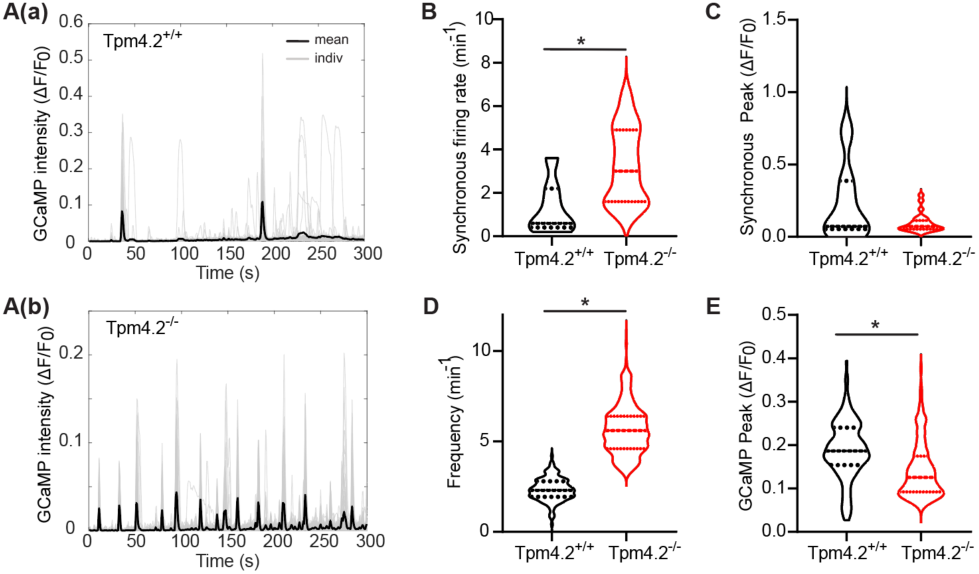
Analysis of network and single neuron Ca^2+^ spikes in Tpm4.2^-/-^ neurons. **A** Representative individual neurons (grey lines) and field of view averaged (black) fluorescent Ca^2+^ signal traces for Tpm4.2^+/+^ and Tpm4.2^-/-^ neurons 20 DIV. **B** Synchronous Ca^2+^ spikes were more frequent in Tpm4.2^-/-^ cultures. **C** The amplitude of synchronous spikes did not differ. **D** The Ca^2+^ spikes in individual Tpm4.2^-/-^ neurons were more frequent. **E** These Ca^2+^ spikes had a smaller amplitude than those observed in Tpm4.2^+/+^ neurons. Data are represented as min-max violin plots with median and quartiles indicated by solid and dashed lines respectively. * indicates p < 0.05.

#### 2.3.2 Electrophysiological Analysis

##### 2.3.2.1 mEPSCs in dissociated cultures of primary Tpm4.2^+/+^ and Tpm4.2^-/-^ neurons

To investigate whether depleting Tpm4.2 would reduce F-actin stability and therefore affect synaptic function, we recorded mEPSCs from cultured neurons, prepared from embryos of Tpm4.2*^-/-^* mice, and Tpm4.2^+/+^ controls (**Figure 4A)**. Tpm4.2*^-/-^* cells had significantly reduced mEPSC frequency (**Figure 4B**; Tpm4.2^+/+^: 8.20 ± 1.26 Hz; Tpm4.2*^-/-^* 3.32 ± 0.66 Hz; Mann-Whitney test p = 0.002) and amplitude (**Figure 4F**; Tpm4.2^+/+^: 33.79 ± 2.69 pA; Tpm4.2*^-/-^*: 22.56 ± 1.53 pA; unpaired t-test t = 3.57, p = 0.001). Tpm4.2*^-/-^* mEPSCs had increased rise time (**Figure 4G**; Tpm4.2^+/+^: 0.58 ± 0.04 ms; Tpm4.2*^-/-^*: 0.70 ± 0.03 ms; unpaired t-test p = 0.02) and decay (**Figure 4H**; Tpm4.2^+/+^: 3.85 ± 0.32; Tpm4.2*^-/-^*: 5.3 ± 0.29; unpaired t-test p = 0.002). We also noted that Tpm4.2*^-/-^*neurons had a higher membrane resistance (**Figure 4D**; Tpm4.2^+/+^: 226.1 ± 27.88; Tpm4.2*^-/-^*: 337 ± 39.77; Mann-Whitney test p = 0.03) and lower membrane capacitance (**Figure 4E**; Tpm4.2^+/+^: 105 ± 4.11 pF; Tpm4.2*^-/-^*: 94.45 ± 2.62 pF; unpaired t-test t = 2.14, p = 0.04). Relative amplitude frequency was also significantly reduced (**Figure 4I)** and relative inter-event interval frequency was significantly increase in Tpm4.2^-/-^ neurons (**Figure 4J**).

**Figure 4.**
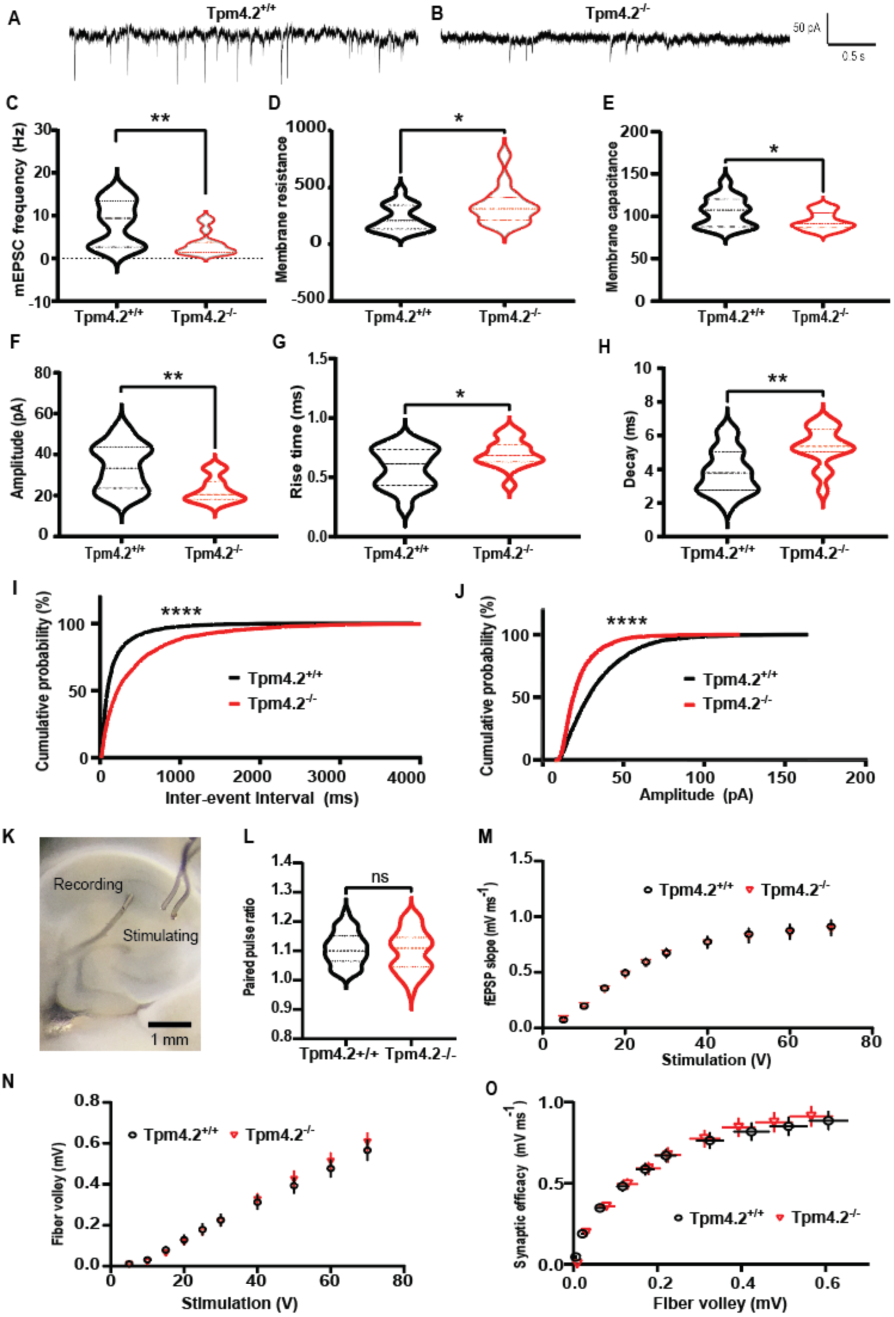
Electrophysiology analysis of Tpm4.2^-/-^ neurons and acute brain slices. **A** Example recording traces for Tpm4.2^+/+^ and **B** Tpm4.2^-/-^ **C** Mean mEPSC frequency was significantly decreased in Tpm4.2^-/-^ neurons. **D** Tpm4.2^-/-^ neurons had a higher membrane resistance and **E** lower membrane capacitance than Tpm4.2^+/+^ neu- rons. **F** Mean mEPSC amplitude was significantly decreased in Tpm4.2^-/-^ neurons. **G** Mean mEPSC rise time and **H** decay were increased in Tpm4.2^-/-^ neurons. **I** Cumulative probability histograms of mEPSC inter-event interval and **J** amplitude; 200 events sampled from each cell were significantly different in Tpm4.2^-/-^ neurons. n = 17 cells from 3 separate culture preparations. K-O Measuring basal synaptic activity in Tpm4.2^-/-^ brain slices **K** Locations of recording and stimulating electrodes in CA1 hippocampus. **L** Paired pulse ratio, a measure of synaptic release probability. **M** IO curve of fEPSP slope. **N** IO curve of fiber volley. **O** fEPSP slope plotted against fiber volley to show synaptic efficacy. Tpm4.2^+/+^ n = 28 slices from 19 mice, Tpm4.2^-/-^ n = 31 slices from 19 mice. Data are represented as min-max violin plots with median and quartiles indicated by solid and dashed lines respectively. Significance was determined as * p < 0.05; ** p < 0.01; **** p < 0.0001. ns = not significant

##### 2.3.2.2 Field excitatory postsynaptic potentials (fPESPs) in Tpm4.2^+/+^ and Tpm4.2^-/-^ brain slices

###### 2.3.2.2.1 The effect of Tpm4.2 knock-out on basal synaptic transmission

As the depletion of Tpm4.2 caused a significant reduction in mEPSC amplitude and frequency in dissociated primary neuron cultures, we examined the effect in acute brain slices prepared from Tpm4.2^-/-^ mice (**Figure 4K**). An input-output (IO) curve was constructed to assess basal function. There was no significant difference between groups in baseline paired pulse ratio (PPR) (**Figure 4L**, unpaired t-test p = 0.57), fEPSP slope (**Figure 4M**; two-way repeated measures ANOVA interaction p = 0.96, genotype p = 0.87), fiber volley (**Figure 4N**; two-way repeated measures ANOVA interaction p = 0.79, genotype p = 0.85), or synaptic efficacy (**Figure 4O**).

###### 2.3.2.2.2 The effect of Tpm4.2 knock-out on LTP

The effect of Tpm4.2 deletion on synaptic plasticity was assessed, using extracellular field potentials in brain slices. After fEPSP amplitude stabilization and induction of LTP (**Figure 5A**), there was no change in the magnitude of LTP induced between groups, either at the early or late stages (**Figure 5B**; two-way ANOVA interaction p = 0.52, genotype p = 0.69). The average level of stimulation chosen to run the protocol based on the IO curve was similar for both groups (**Figure 5C-D**; Tpm4.2^+/+^ = 12.93 ± 0.35 V; Tpm4.2^-/-^ = 12.63 ± 0.44 V; p = 0.62). PPR was measured throughout to check for presynaptic changes. We observed no significant differences in PPR at any point in the experiment (**Figure 5E**; two-way RM ANOVA interaction p = 0.33, genotype p = 0.07).

**Figure 5.**
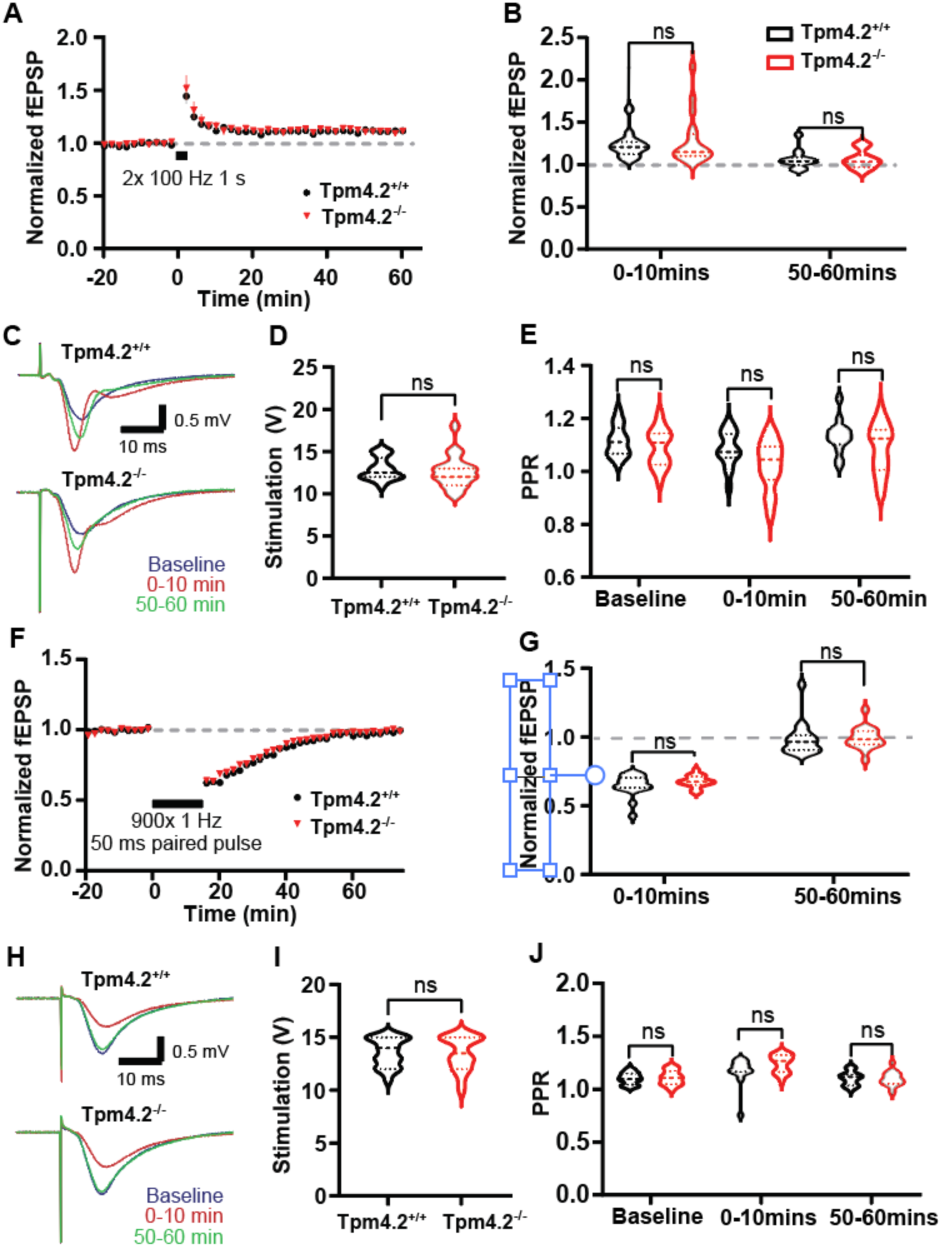
The effect of Tpm4.2 knock-out on LTP (A-E) and LTD (F-J) in acute brain slices. **A** Plot of fEPSP over course of experiment, with each point representing the average from 2 minutes of recording for LTP. **B** Mean normalized fEPSP at first and last 10 minutes of recording after inducing LTP were not significantly different between Tpm4.2^+/+^ (black) and Tpm4.2^-/-^ (red) cultures. **C** Example waveforms from baseline (blue), first 10 minutes after low frequency stimulation (red) and last 10 minutes of recording (green) for LTD. **D** Stimulation intensities used for LTP. **E** PPR at baseline and first and last 10 minutes after high frequency stimulation was not significantly different between groups for LTP. **F** Plot of fEPSP over course of experiment, with each point representing the average from 2 minutes of recording for LTD. **G** Mean normalized fEPSP at first and last 10 minutes of recording after inducing LTD were not significantly different between Tpm4.2^+/+^ (black) and Tpm4.2^-/-^ (red) cultures **H** Example waveforms from baseline (blue), first 10 minutes after low frequency stimulation (red) and last 10 minutes of recording (green) for LTD. **I** Stimulation intensities used for LTD were the same as used for those in LTP (**D**) experiments. **J** PPR at baseline and first and last 10 minutes after stimulation was not significantly different between groups for LTD. Tpm4.2^+/+^ n = 19 slices from 11 mice, Tpm4.2^-/-^ n = 18 slices from 9 mice.

###### 2.3.2.2.3 The effect of Tpm4.2 knock-out on LTD

A stimulation level evoking approximately half of the maximal response was chosen. A stable baseline of at least 20 minutes was recorded, before 15 minutes of a 1 Hz paired pulse to induce LTD (**Figure 5F)**. No significant differences in the level of depression either at the early or late stage or recording were identified (**Figure 5G**; two-way repeated measures ANOVA interaction p = 0.58, genotype p = 0.42). The stimulation level used for the experiments did not differ between the two groups (**Figure 5H-I**; Tpm4.2^+/+^ = 13.58 ± 0.31 V; Tpm4.2^-/-^ = 13.5 ± 0.38 V; unpaired t-test t = 0.16, p = 0.87). The PPR was also not significantly altered (**Figure 5J**; two-way repeated measures ANOVA interaction p = 0.07, genotype p = 0.25).

### 2.4 Neurite complexity of Tpm4.2 knock-out neurons

To investigate whether Tpm4.2 knock-out affects neuronal morphology, morphometric analysis was conducted to quantify neurite outgrowth in Tpm4.2^-/-^ compared with Tpm4.2^+/+^ neurons. Tpm4.2^-/-^ axons were significantly longer than axons from Tpm4.2^+/+^ neurons (37.91% increase, p < 0.0001, **Figure 6A**). There was no significant difference in the number of axon primary branches (p > 0.05, **Figure 6B)** or axon secondary branches (p > 0.05, **Figure 6C)** between Tpm4.2^+/+^ and Tpm4.2^-/-^ neurons.

**Figure 6.**
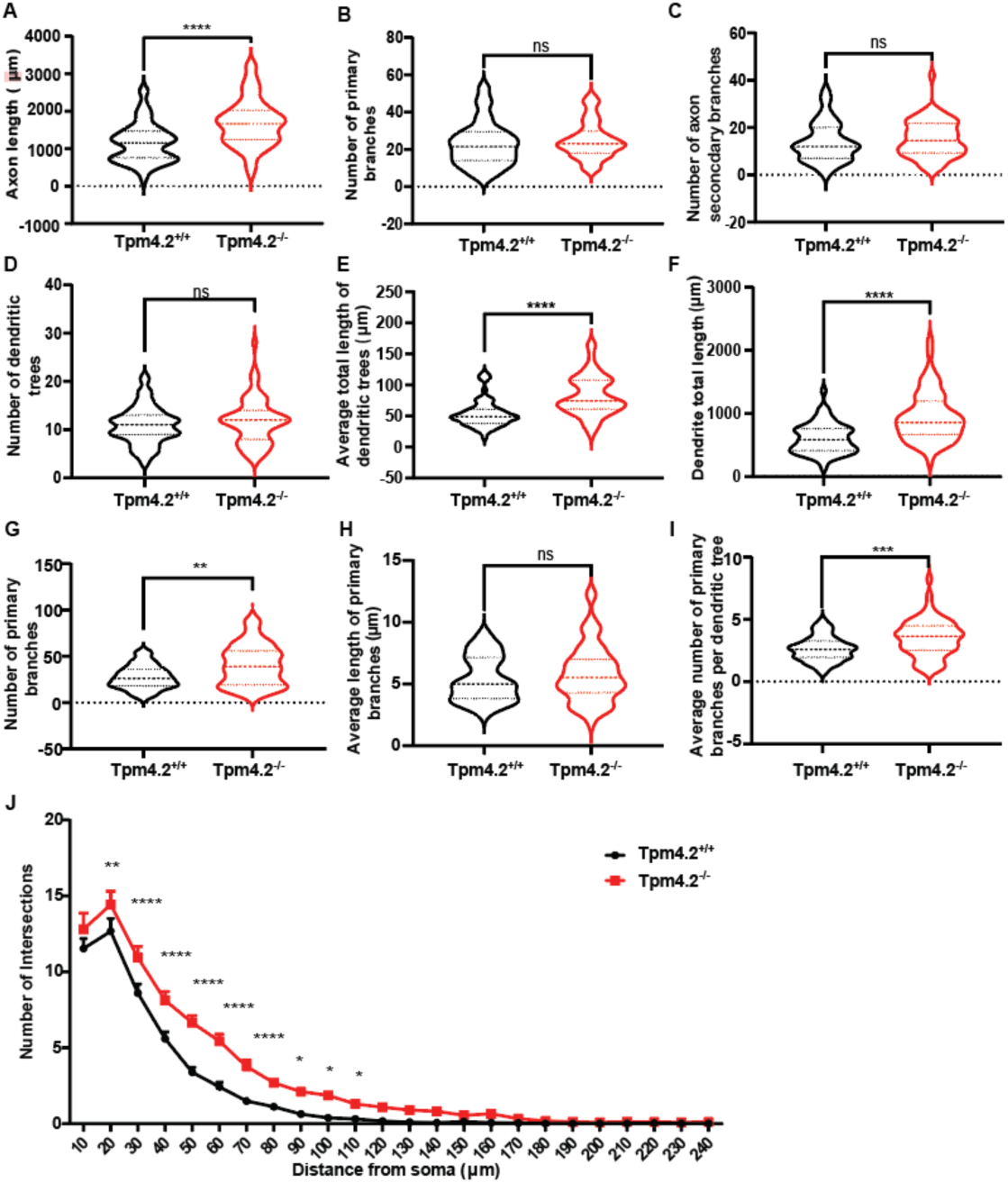
Neurite and spine changes in Tpm4.2^-/-^ neurons. **A**-**C** Differences in axonal complexity and **D**-**I** dendritic complexity between Tpm4.2^+/+^ and Tpm4.2^-/-^ neurons. **A-I** Data are depicted in min-max violin plots with median and quartiles indicated by solid and dashed lines respectively. **J** Sholl analysis of Tpm4.2^-/-^ neurons compared with Tpm4.2^+/+^. Tpm4.2^+/+^ (black) n = 51 cells from 4 separate culture preparations, Tpm4.2^-/-^ (red) n = 57 neurons from 3 culture preparations. Significance was calculated using Mann-Whitney U test. ns = not significant, ** p < 0.01; *** p < 0.005; **** p < 0.0001.

There was no significant difference in the number of dendritic trees in Tpm4.2^-/-^ neurons (p > 0.05, **Figure 6D)**. However, there was a significant increase in the length and complexity of Tpm4.2^-/-^ dendrites compared with Tpm4.2^+/+^ neurons. Tpm4.2^-/-^ neurons had a higher mean total length of dendritic trees (52.62% increase, p < 0.0001; **Figure 6E**), dendrite total length (56.68% increase, p < 0.0001, **Figure 6F)** and number of primary branches (47.86% increase, p = 0.0012, **Figure 6G).** The mean length of primary branches was unchanged (**Figure 6H)**. Tpm4.2^-/-^ neurons had a higher mean number of primary branches per dendritic tree (33.02% increase, p = 0.0004, **Figure 6I**. Sholl analysis identified that this increased dendritic complexity occurred 20-90 μm from the soma (p < 0.0001, **Figure 6J**)

### 2.5 Spine morphology in Tpm4.2^-/-^ neurons

There was no significant difference in dendritic spine density between the Tpm4.2^-/-^ and Tpm4.2^+/+^ neurons (**Figure 7A-B**; Tpm4.2^+/+^: 0.87 ± 0.03 μm -1; Tpm4.2^-/-:^ 0.8 ± 0.03 μm-1; unpaired t-test t = 2.51, p = 0.25). There was also no significant difference in the mean length (**Figure 7C**; Tpm4.2^+/+^: 0.71 ± 0.02 μm; Tpm4.2^-/-^: 0.74 ± 0.02 μm; unpaired t-test t = 2.06, p = 0.29) or width (**Figure 7D**; Tpm4.2^+/+^: 0.39 ± 0.01 μm; Tpm4.2^-/-^: 0.37 ± 0.01 μm; unpaired t-test t = 1.32, p = 0.19) of the dendritic spines. However, the cumulative probability histograms showed a shift towards longer (**Figure 7E**; Kolmogorov-Smirnov p = 0.007) and thinner (**Figure 7F**; Kolmogo-rov-Smirnov p < 0.0001). Additionally, there was a difference in distribution of dendritic spine types (**Figure S4**; two-way ANOVA interaction p = 0.03). Tukey post-hoc multiple comparisons however did not detect any significant differences between groups (p > 0.05 for all), however there was a trend towards a decrease in stubby dendritic spines in Tpm4.2^-/-^ neurons compared with Tpm4.2^+/+^ (**Figure S4;** 0.1661 vs 0.1212 ± 0.018, 36.64% reduction; p = 0.067).

**Figure 7.**
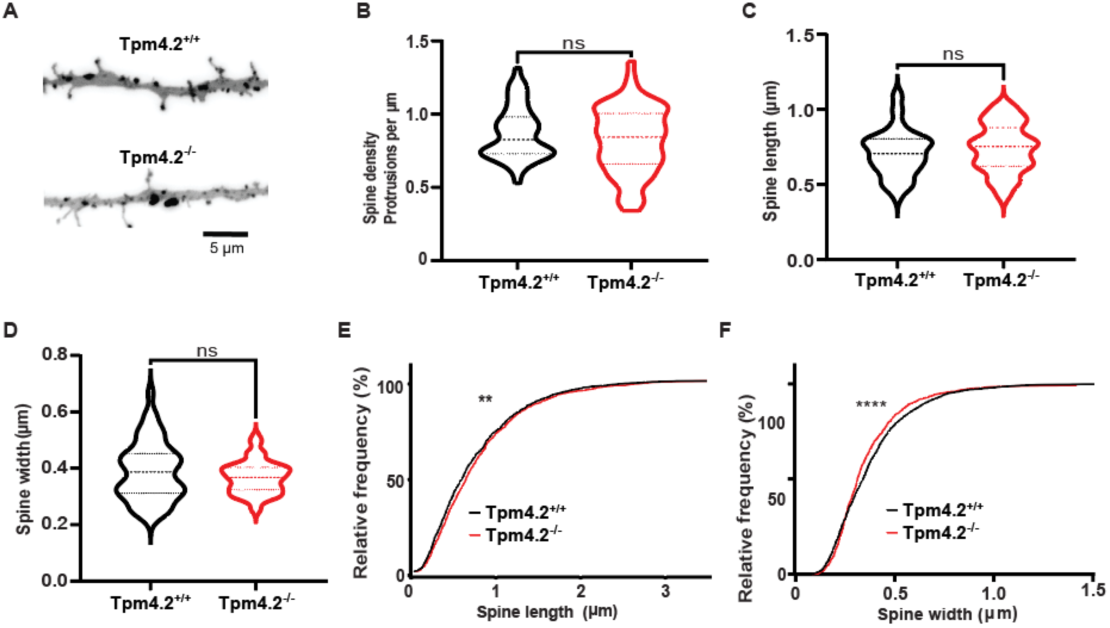
Spine analysis of Tpm4.2^-/-^ neurons. **A** Representative inverted dendrite images flattened from z-stack. **B** Spine density, **C** mean spine length and **D** mean spine widths are similar between Tpm4.2^+/+^ and Tpm4.2^-/-^ neurons. Cumulative frequency histograms of **E** spine length and **F** spine width with Tpm4.2^+/+^ depicted with a black line and Tpm4.2^-/-^ represented by a red line. Tpm4.2^+/+^ n = 51 cells from 4 separate culture preparations, Tpm4.2^-/-^ n = 57 neurons from 3 culture preparations. Significance was calculated using Mann-Whitney U test. ns = not significant, ** p < 0.01; **** p < 0.0001.

**Figure S4.**
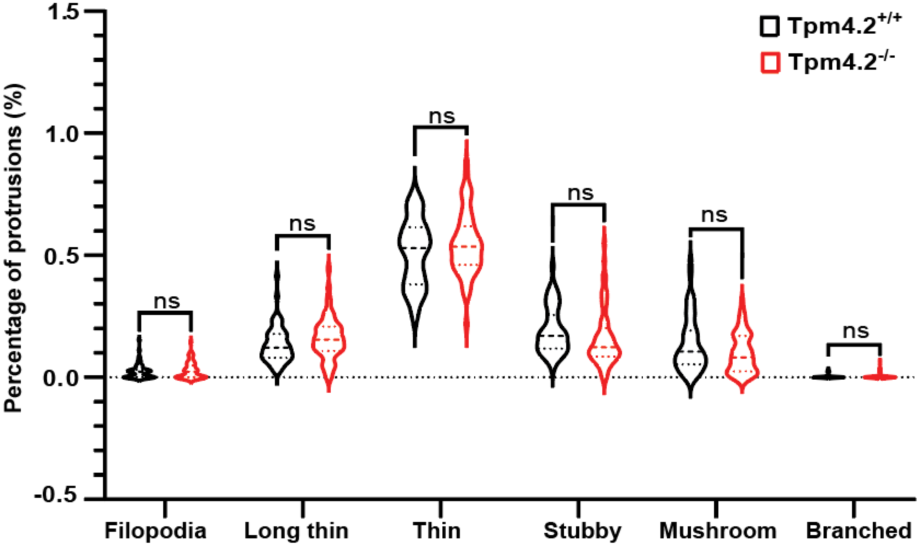
Spine categories for Tpm4.2^-/-^ neurons. There were no significant differences in spine categories between Tpm4.2^+/+^ and Tpm4.2^-/-^ neurons. There was a trend towards a decrease in stubby dendritic spines in Tpm4.2^-/-^ neurons p = 0.06. ns = not significant.

### 2.6 Mouse behavioral testing

We performed a battery of behavioral tests on Tpm4.2^+/+^ and Tpm4.2^-/-^ littermates that assessed anxiety, learning and socialization to test the effect of Tpm4.2 on these cognitive aspects.

#### 2.6.1. Open field (OF)

Three-way repeated measures ANOVA revealed a significant main effect of ‘time’ on the OF distance travelled (F(5,205) = 100.5, p < 0.001, **Figure 8A**). All animals exhibited a decline in the distance travelled across time, with females displaying greater locomotion than males (main effect of ‘sex’: F(1,41) = 5.7, p = 0.022, **Figure 8A**). A sex effect was also demonstrated in the cumulative distance travelled over the 30-minute protocol, (F(1,41) = 5.7, p = 0.022, **Figure 8B**). No discernible ‘genotype’ effect on distance travelled was detected (F(1,41) = 0.625, p = 0.434; **Figure 8C**). The rearing frequency was similar between sexes (F(1,41) = 1.5, p = 0.223, **Figure 8D**) and genotypes (F(1,41) = 1.5, p = 0.231, **Figure 8D**).

**Figure 8.**
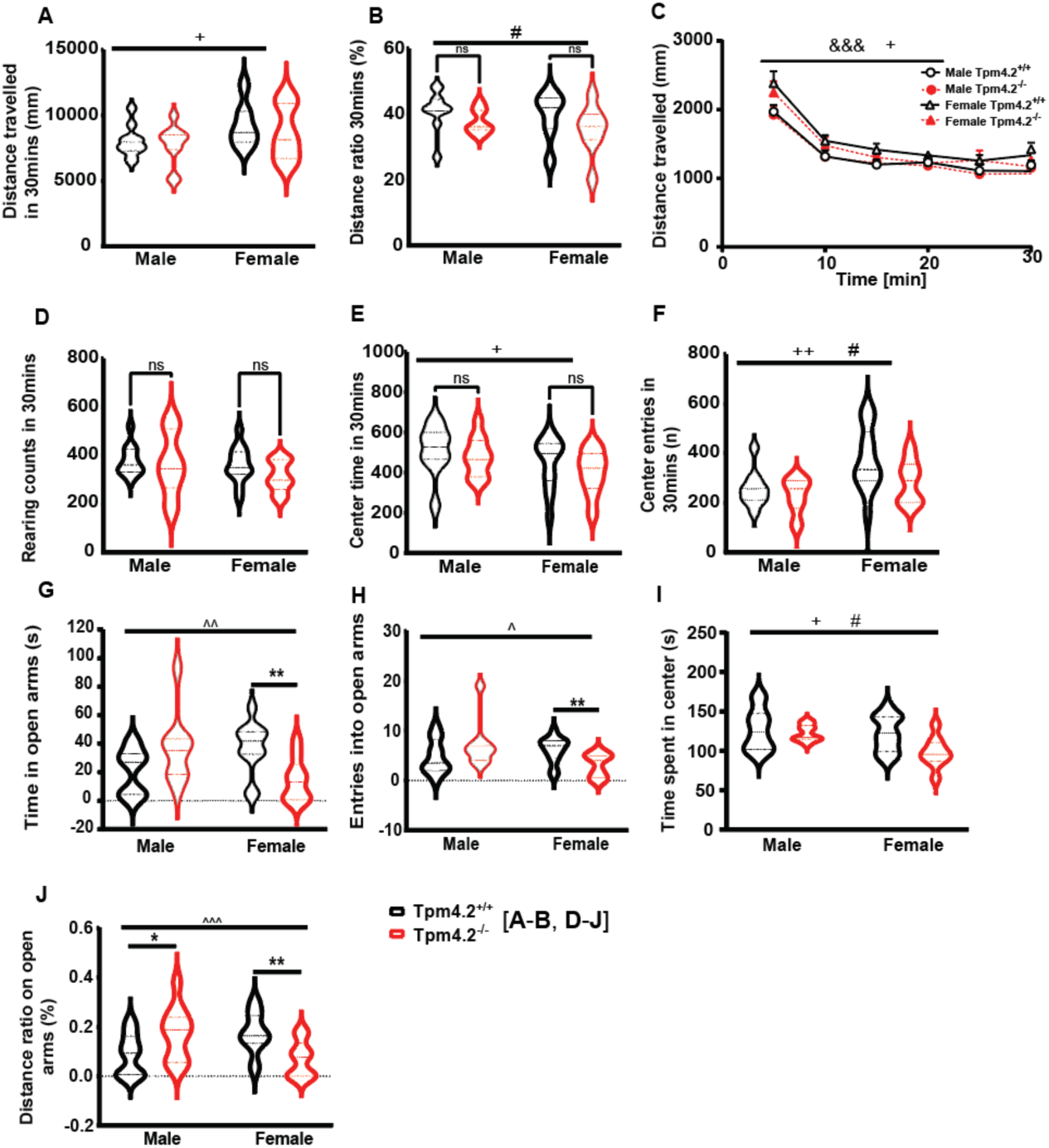
Assessing locomotion and anxiety-like behaviors in Tpm4.2^-/-^ mice. **a-g)** Open field behaviors – Locomotion, exploration and anxiety. **a)** Distance travelled (mm) across 5-minute blocks **(b)** total distance travelled **(c)** frequency of rearing (i.e. vertical activity) **(d)** Distance travelled in the center of the OF divided by total distance, **(e)** center time and **(f)** entries. **(h-j)** Elevated plus maze behaviors – Anxiety with **(h)** Entries in the open arms of the EPM and **(b)** time spent in the EPM center and **(j)** ratio of distance travelled in the center. Data are shown as violin plots of mean ± SEM for male and female Tpm4.2^+/+^ and Tpm4.2^-/-^ littermates. Tpm4.2^+/+^ (black) n = 25 mice, Tpm4.2^-/-^ (red) n = 20 mice. Three-way repeated measured or two-way ANOVA were used, ‘time’, ‘sex’, ‘genotype’ and ‘interaction’ effects are indicated by &&& (p < 0.001), + (p < 0.05 - ++ p < 0.01) and # (p < 0.05), ^ (p < 0.05 - ^^p < 0.01 - ^^^p < 0.001), respectively. Unpaired t-test significances are indicated by * (p < 0.05 - **p < 0.01).

When assessing anxiety behaviors, females spent less time (F(1,41) = 4.6, p = 0.037, **Figure 8E**) and had fewer entries (F(1,41) = 10.4, p = 0.003, **Figure 8f**) in the open field center compared to their male counterparts. Notably, a main effect of ‘genotype’ was evident for entries (F(1,41) = 4.089, p = 0.049, **Figure 8F**) and distance ratio (F(1,41) = 4.614, p = 0.037, **Figure 8C**), with Tpm4.2^-/-^ mice exhibiting less entries and reduced locomotion in the centre of the open field, indicating heightened anxiety levels in Tpm4.2^-/-^ mice relative to Tpm4.2^+/+^ regardless of sex (no ‘sex’ by ‘genotype’ interaction, p > 0.05). There was a significant interaction in time spend in open arems (p = 0.0012, **Figure 8G**) and post-hoc multiple comparisons identified a significant difference in time spent in open arms for female Tpm4.2^+/+^ compared with female Tpm4.2^-/-^ mice (88.70% reduction, p = 0.0305, **Figure 8G**)

#### 2.6.2 Elevated Plus Maze (EPM)

Behaviors assessed using EPM appeared largely sex-specific, with male Tpm4.2^-/-^ mice demonstrating an anxio-lytic-like phenotype whereas female Tpm4.2^-/-^ mice had higher anxiety levels compared to the respective controls. Two-way ANOVA revealed significant ‘sex’ by ‘genotype’ interactions for entries into open arms (F(1, 41) = 6.712, p = 0.013, **Figure 8H**) time spent on (F(1, 41) = 12.13, p = 0.001, **Figure 8I**) and as well as for the distance ratio exploring the open arms (F(1, 41) = 12.65, p = 0.001, **Figure 8J**). Subsequent post-hoc analysis splitting the data by sex revealed that female Tpm4.2^-/-^ mice exhibited fewer entries (t(20) = 2.811, p = 0.010, **Figure 8H**) and spent less time (t(20) = 3.269, p = 0.003, **Figure 8I**) in the open arms than their respective controls, whereas no genotype differences were observed for these behaviors in male mice. Conversely, Tpm4.2^-/-^ males exhibited increased locomotion ratio in the open arms compared to Tpm4.2^+/+^ control males (t(21) = 2.102, p = 0.047, **Figure 8J**), while female Tpm4.2^-/-^ mice displayed a decreased ratio distance in the open arms (t(20) = 3.037, p = 0.006, **Figure 8J**). Additionally, a main effect of sex (F(1, 41) = 5.817, p = 0.020) and genotype (F(1, 41) = 5.017, p = 0.0301) was observed for the time spent in the center of the maze (**Figure 8I**).

#### 2.6.3 Social Preference Test (SPT)

A three-way ANOVA revealed a significant main effect of side for the nosing time in both the sociability trial (F (1, 35) = 96.07, p < 0.001, **Figure 9A**) and the social novelty preference trial (F (1, 35) = 18.39, p < 0.001, **Figure 9B**). Subsequent paired t-tests revealed that, as expected, all groups exhibited a preference for exploring the A/J mice over the empty chamber in the sociability test (Tpm4.2^+/+^ male t(7) = 3.999, p = 0.0052; Tpm4.2^-/-^ male t(8) = 4.048, p = 0.0037; Tpm4.2^+/+^ female t(12) = 7.987, p < 0.001; Tpm4.2^-/-^ female t(8) = 4.528, p = 0.0019, **Figure 9A**). However, in the social novelty preference test (**Figure 9B**), only males demonstrated a heightened preference for the new animal over the familiar one for both genotypes (Tpm4.2^+/+^ male t(7) = 3.750, p = 0.007; Tpm4.2^-/-^ male t(8) = 2.331, p = 0.048). Female Tpm4.2^+/+^ mice showed a trend towards exploring the new A/J mice more than the familial one (t(12) = 2.050, p = 0.062), whereas female Tpm4.2^-/-^ did not display any preference (t(8) = 0.582, p = 0.576), potentially indicating an impairment in social memory. Similar findings were observed using one-sample t-tests to analyze percentage exploration compared to chance exploration. While all animals showed increased preference for the side with the A/J mouse (p < 0.05 for all, **Figure S5A** for social preference test, on the social novelty test only males preferred the new mice over the familiar ones (p > 0.05 for all, **Figure S5B**). The female Tpm4.2^+/+^ displayed a trend in exploration index (t(12) = 1.183, p = 0.091, **Figure 5B**), while the Tpm4.2^-/-^ females explored both mice equally (t(8) = 0.477, p = 0.645, **Figure S5B**).

**Figure 9.**
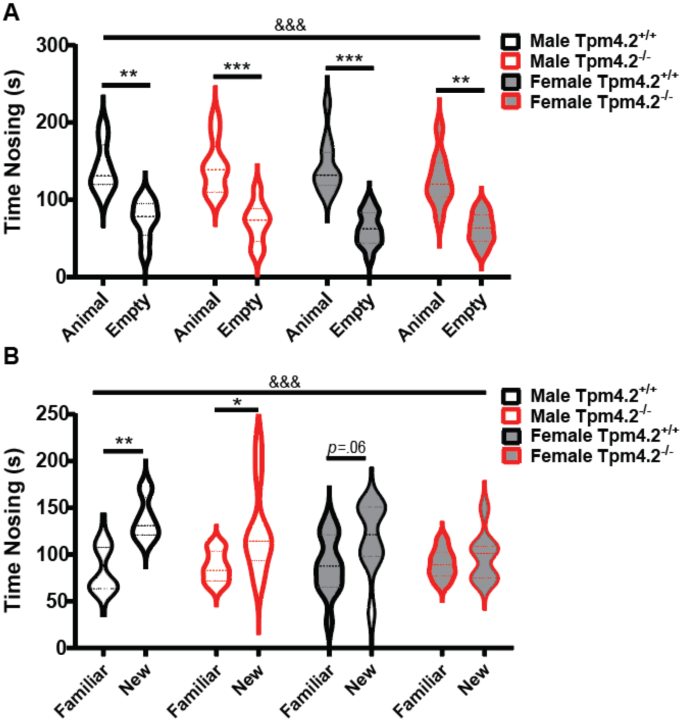
Social Preference Test in Tpm4.2^-/-^ mice. Sociability test behaviors – Social preference and social novelty. Total time spent nosing chambers containing **A** an A/J mouse versus no mouse (i.e. empty chamber) (sociability) or **B** a novel versus a familial A/J mouse (social novelty preference). Data are shown as mean ± SEM for Tpm4.2^+/+^ (black) n = 25 mice, Tpm4.2^-/-^ (red) n = 20 mice. Three-way RM ANOVA effects of ‘side’ are indicated by &&& (p < 0.001). Paired t-test significances are indicated by * (p < 0.05 - ***p < 0.001).

**Figure S5.**
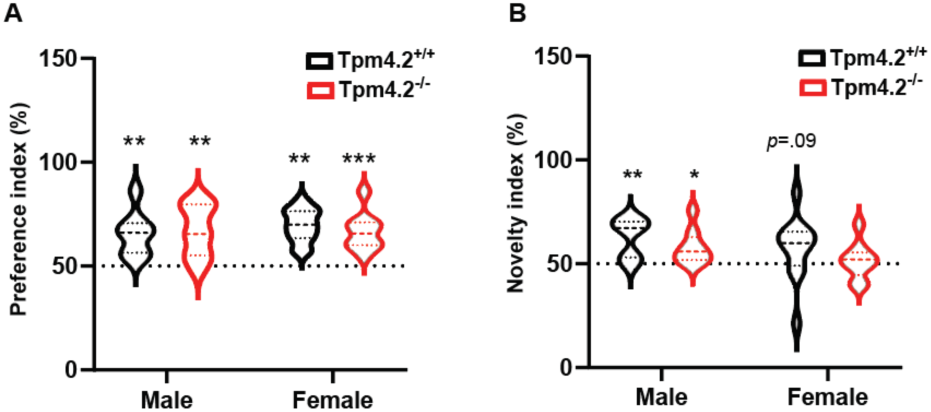
Sociability test behaviours – Social preference and social novelty. Percentage of time spent nosing chambers containing **A** an A/J mouse versus no mouse (i.e. empty) (sociability) or **B** a novel versus a familial A/J mouse (social novelty preference), represented by the exploration index (social preference test: mouse x 100/ mouse + empty or social novelty test: new mouse x 100/ new + familiar). Data are shown as mean ± SEM for male and female Tpm4.2^+/+^ (black) n = 25 mice, Tpm4.2^-/-^ (red) n = 20 mice. Three-way repeated measured or two-way ANOVA were used, ‘time’, effects are indicated by &&& (p < 0.001), One sample t-test results against chance levels are indicated by * (p < 0.05 - ***p < 0.001).

#### 2.6.4 Novel Object Recognition Test (NORT)

A three-way repeated measures ANOVA detected a significant main effect of ‘object’ (F(1,38) = 8.129, p = 0.007] and ‘genotype’ (F(1,38) = 6.466, p = 0.015), with the latter indicating that Tpm4.2^-/-^ mice exhibited lower levels of exploration compared to Tpm4.2^+/+^ across sex (**Figure 10**). Additionally, a significant ‘object’ by ‘sex’ by ‘genotype’ triple interaction was observed (F(1,38) = 4.459, p = 0.041, **Figure 10A**), with subsequent paired t-tests elucidating an increased exploration of the new object by Tpm4.2^-/-^ males([t(10) = 3.407, p = 0.006, Fig. 4A), which was also a trend for females Tpm4.2^+/+^ mice (t(11) = 2.035, p = 0.066). Male Tpm4.2^+/+^ controls (t(8) = 1.478, p = 0.177, **Figure 10**) and Tpm4.2^-/-^ females (t(9) = 0.704, p = 0.499, **Figure 10**) failed to show a preference for the novel object. To analyze percentage exploration compared to chance levels with one-sample t-tests we observed a similar finding (**Figure S6**). Male Tpm4.2^-/-^ mice had a preference for the novel object (t(10) = 3.644, p = 0.0045), whereas their respective Tpm4.2^+/+^ controls exhibit no statistically significant difference (t(8) = 1.779, p = 0.113). Again, the female Tpm4.2^+/+^ displayed a trend in nosing index (t(11) = 2.176, p = 0.052), while the Tpm4.2^-/-^ females explored both objects equally (t(9) = 0.216, p = 0.833).

**Figure 10.**
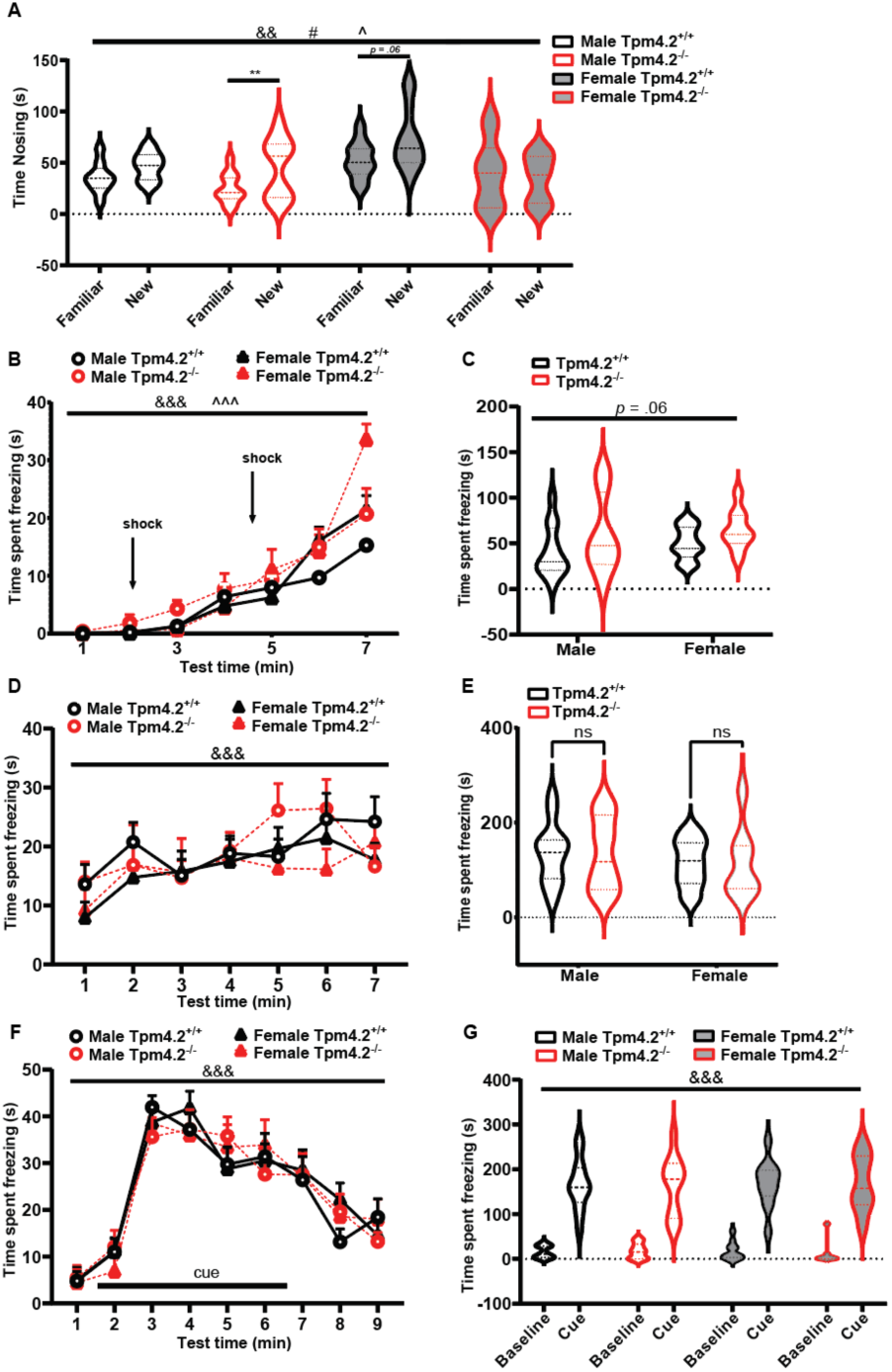
Assessing learning and memory in Tpm4.2^-/-^ mice. **a)** Novel object recognition behaviors - recognition memory. Time nosing the familiar or the new object. **(b-f)** Contextual and Cue fear conditioning. Freezing in the fear conditioning test – fear-associated memory. Time spent freezing **(b,c)** during conditioning, **(d, e)** during context testing or **(f,g)** during cue testing for either **(b,d)** every 1-minute block, **(c,e,f)** as a total during test duration or **(f)** averaged per minute for the first 2 minutes (cue off) compared to the following 5 min (cue on) of the cue testing. Data are shown as mean ± SEM for male and female Tpm4.2^+/+^ (black) n = 25 mice, Tpm4.2^-/-^ (red) n = 20 mice littermates. Three-way ANOVA effects of ‘object’, ‘genotype’ and interaction thereof are indicated by && (p < 0.01), &&& (p < 0.001), # (p < 0.05) and ^ (p < 0.05) ^^^ (p < 0.001), respectively. Paired t-test results are indicated by ns = not significant ** (p < 0.01) or exact p-value.

**Figure S6.**
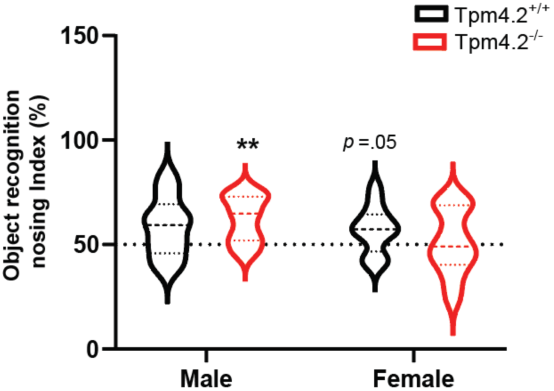
Novel object recognition behaviours - recognition memory. Nosing index, represented by the time exploring the new object as a percentage of total object exploration (%). Data are shown as mean ± SEM for male and female wild type-like Tpm4.2^+/+^ and Tpm4.2^-/-^ littermates. One sample t-test results against chance levels are indicated by ** (p < 0.01).

#### 2.6.5 Contextual and Cue Fear Conditioning (CFC)

During the conditioning phase, a three-way repeated measures ANOVA revealed a significant main effect of ‘time’ (F(6, 246) = 93.80, p < 0.001, **Figure 10B**), indicating an overall increase in freezing post foot-shock across all animals. While there were no discernible effects of ‘sex’ (F(1, 41) = 0.696, p = 0.409, **Figure 10B)** or ‘genotype’ (trend only: F(1,41) = 3.621, p = 0.064, **Figure 10B**), interactions of ‘time’ with ‘sex’ (F(6, 246) = 5.736, p < 0.001, **Figure 10B**) and with ‘genotype’ (F(6, 246) = 3.142, p = 0.005, **Figure 10B**) were evident. However, further examination using a two-way ANOVA split by ‘time’ disclosed effects of ‘genotype’ and ‘sex’ only at minute 7 (sex: F(1, 42) = 7.069, p = 0.011 – genotype: F(1, 42) = 6.466, p = 0.014, **Figure 10B**). Tpm4.2^-/-^ mice exhibited heightened freezing in the final minute of test following two foot shocks compared to Tpm4.2^+/+^ controls (p < 0.05), and Tpm4.2^-/-^ females also displayed increased freezing compared to Tpm4.2^-/-^ males (p < 0.05). Total freezing time analysis suggested a trend for a ‘genotype’ effect (F(1, 41) = 3.621, p = 0.064, **Figure 10C**), yet no sex effect was evident (F(1, 41) = 0.6955, p = 0.4091, **Figure 10C**).

In the context test, freezing increased over time as expected (‘time’: F(6, 246) = 7.301, p < 0.001, **Figure 10D**). No effects of ‘sex’ (F(1, 41) = 1.132, p = 0.293, **Figure 10D**) or ‘genotype’ (F(1, 41) = 0.006, p = 0.937, **Figure 10D**) and no interactions with ‘time’ (all p’s > 0.05) were discerned. Two-way ANOVA for total freezing time during the context test revealed no significant differences either (p > 0.05 for all, **Figure 10E**).

A three-way repeated measures ANOVA for the cue trial demonstrated an overall effect of ‘time’ (F(8, 328) = 70.95, p < 0.001, **Figure 10F**), with all animals exhibiting increased freezing levels in response to the cue. No effects of ‘sex’ (F(1, 41) = 0.038, p = 0.845, **Figure 10F**) or ‘genotype’ (F(1, 41) = 0.008, p = 0.931, **Figure 10F**) were observed. Similar non-significant results were found for total freezing time during cue presentation (3rd to 7th minute; data not shown). An overall effect of ’cue x baseline’ (average freezing per minute during baseline versus cue freezing, **Figure 10G**) was identified in the three-way repeated measures ANOVA (F(1, 42) = 357.7, p < 0.001, **Figure 10G**) confirming that the freezing response increases at cue presentation, while no effects of ‘sex’ (F(1, 42) = 0.0208, p = 0.885, **Figure 10G**) or ‘genotype’ (F(1, 42) = 0.032, p = 0.858, **Figure 10G**) were evident or interfered with the overall freezing response to the cue (no interactions with ‘time’, p > 0.05 for all).

#### 3.6.3 Prepulse Inhibition (PPI)

In examining the startle response, a three-way repeated measures ANOVA yielded a significant main effect of ‘startle intensity’ (F(2, 82) = 122.9, p < 0.001, **Figure S7A**), demonstrating that the startle response increased in tandem with the intensity across all groups. No significant effects were observed for sex (F(1, 41) = 3.408, p = 0.0721, **Figure S7A**) or genotype (F(1, 41) = 2.797, p = 0.102, **Figure S7A**) and no interactions with ‘startle intensity’ were evident either (all p’s > 0.05). Similarly, analysis of habituation to the startle over time revealed an overall effect for the startle pulse blocks (F(2, 82) = 17.25, p < 0.001, **Figure S7B**), indicating habituation across all groups. No significant differences were found for sex (F(1, 41) = 2.487, p = 0.122, **Figure S7B**) or genotype (F(1, 41) = 2.27, p = 0.139, **Figure S7B**). Finally, when analyzing the prepulse inhibition response, all animals demonstrated an elevated PPI response to increasing prepulse intensities (F(2, 82) = 347.2, p < 0.001, **Figure S7C**) regardless of experimental condition (all other p’s > 0.05).

**Figure S7.**
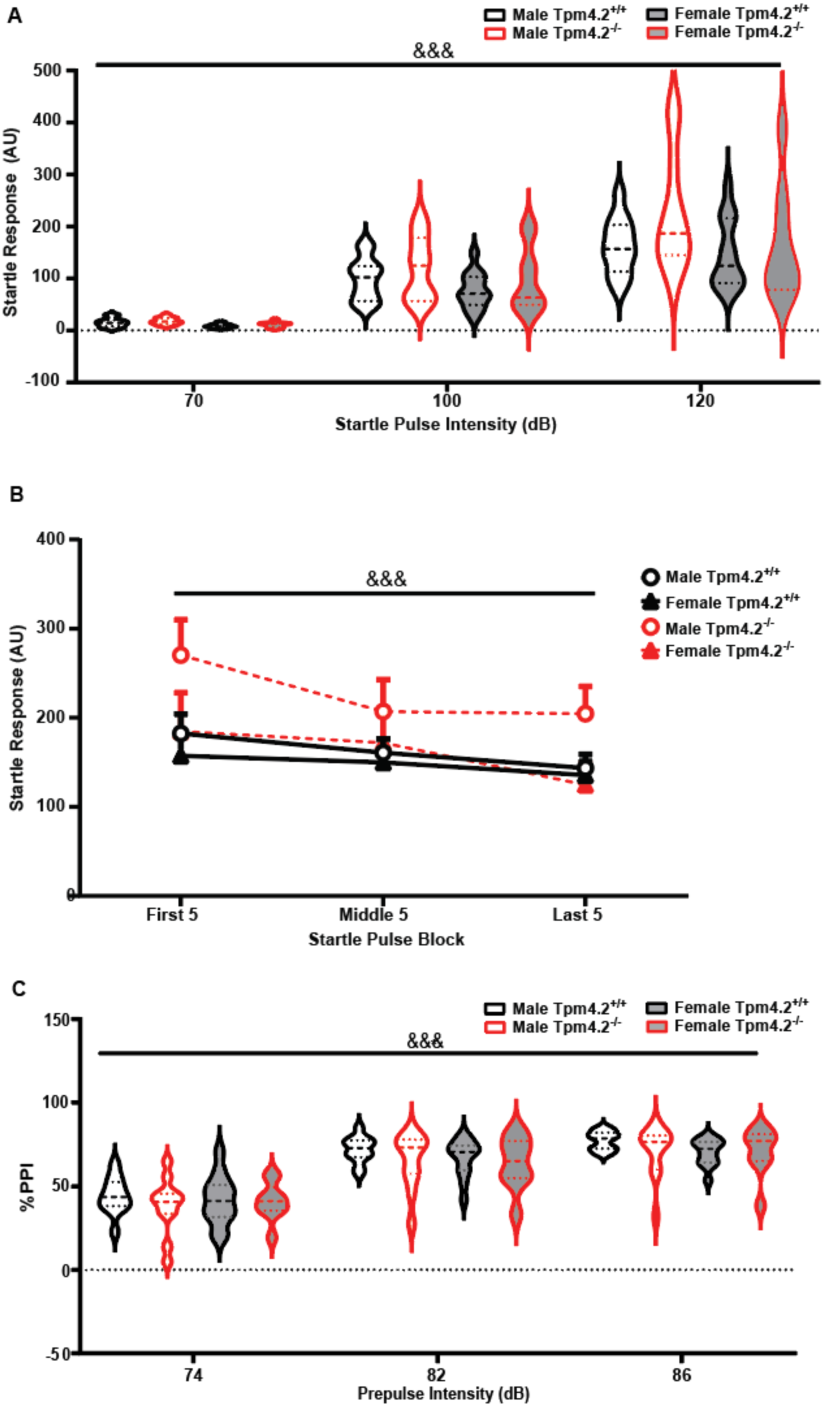
Prepulse Inhibition test in Tpm4.2^-/-^ mice. **A** Average startle response to different pulse intensities. **B** Startle habituation separated by 5-minute blocks. **c)** Average percentage of inhibition by different prepulse intensities. Data are shown as mean ± SEM for male and female Tpm4.2^+/+^ (black) n = 25 mice, Tpm4.2^-/-^ (red) n = 20 mice littermates.

## 3. Discussion

The current study examined Tpm4.2 depletion-dependent behavioral phenotypes and cellular functions of Tpm4.2 in developing and mature neurons, using cellular assays in primary hippocampal neuron cultures and acute brain slices and a battery of cognitive test regimes. Here, we confirm previous reports [10] that Tpm4.2 is expressed throughout the brain and identified a role for Tpm4.2 in aspects of neuronal signaling, with increases in synchronous firing and firing frequency yet impaired neuronal signaling strength, in Tpm4.2 knock-out neurons. Tpm4.2 knock-out mice exhibited a mild behavioral phenotype of anxiety which was predominantly sex-dependent, and subtle reductions in social memory which was more prominent in females. Dissociated neuronal cultures of Tpm4.2-depleted neurons demonstrated increased dendritic complexity, impaired receptor internalization, and increased neuronal network connectivity, however more complex brain slice cultures exhibited no significant changes in basal activity or alterations in either LTP or LTD. These data indicate a physiological role of Tpm4.2 in aspects of neuronal signaling, neurite growth regulation and development, with no discernable effect on overall synaptic plasticity.

Using primary hippocampal neurons as neuronal model, we confirmed a previously suggested role of Tpm4.2 in neurite outgrowth, providing a physiological function for the high expression of Tpm4.2 levels in dendritic growth cones of immature neurons [10], During maturation, Tpm4.2 expression becomes more localized to the post-synapse [12], where we demonstrate a role for Tpm4.2 in signaling frequency, strength and AMPAR recycling at the post-synapse. As Tpm forms co-polymers with F-actin, these data suggest specific roles for the corresponding F-actin subpopulations in normal neuronal health and signaling.

To determine whether Tpm4.2 is involved in cellular processes of learning and memory through long-term potentiation (LTP) and long-term depression (LTD), we analyzed receptor recycling processes, calcium signaling, and mEPSCs in dissociated primary hippocampal neurons as a neuronal model, and fEPSPs in acute slices to provide a more complex biological system more representative of the human brain. We observed a reduced internalization of the GluA1 subunit component of AMPA receptors in Tpm4.2^-/-^ neurons, indicating the potential for overstimulation and excitotoxicity in Tpm4.2-depleted neurons. Selective modifications of post-synaptic AMPA receptors play a key role in the cellular mechanism for learning and memory – LTP [14]. The regulation of AMPAR is essential as it mediates most of the fast excitatory neurotransmission, and the abundance of AMPAR at the surface of excitatory synapses dictates the strength of responses to excitatory stimulation [15]. The AMPAR subunit GluA1 mediates activity-dependent changes in excitatory synaptic transmission between neurons [16]. Internalization of receptors following stimulation is a necessary process to prevent overstimulation. In neurons lacking Tpm4.2 however, GluA1 receptor subunit internalization was impaired, indicating a physiological function of Tpm4.2 in receptor recycling processes, and therefore in mediating aspects of neuronal excitation. Diminished GluA1 internalization and the resulting increase in surface GluA1 receptors can have substantial effects on neuronal excitability and is proposed to contribute to the increased neuronal firing frequency observed in live imaging of calcium levels in Tpm4.2^-/-^ neurons in the current study.

Despite evidence that the actin cytoskeleton plays a key role in processes of endocytosis, including endosomal retrieval and recycling [17], the precise role of actin in AMPAR trafficking is not fully understood. As F-actin is abundant in dendritic spines, it is suggested to play a physiological role in regulating various aspects of the post-synapse – including receptor trafficking. Actin depolymerizing drugs such as latrunculin reduce GluA1-containing AMPARs in dendritic spines [18], and reduced surface expression at synapses [19], while jasplakinolide, an F-actin stabilizing drug prevents AMPAR internalization [20]. In addition, there are reports of AMPAR pools that are regulated by actin processes, and others that are unaffected, suggesting distinct actin subpopulations, some of which affect AMPAR recycling processes, and some that are not involved in this process. As Tpms form co-polymers with F-actin and have previously been suggested to play a role in bulk endocytosis [21], we hypothesized that Tpms are involved in processes of receptor internalization in an isoform-specific manner. Here, we report a role for Tpm4.2 and its associated F-actin subpopulation in AMPAR recycling with impaired GluA1 internalization in Tpm4.2^-/-^ neurons. The concurrent increase in abundance of excitatory surface receptors ultimately increases the likelihood of further activation and hyperexcitability in Tpm4.2^-/-^ neurons.

Dissociated Tpm4.2^-/-^ neuron cultures showed increases in both synchronous and single cell firing frequency when compared with Tpm4.2^+/+^ neurons. However, the calcium response was diminished. The reduced calcium response may reflect a reduction in the number of APs generated within the underlying calcium spike or differences in ion-channel activity. The reduced mEPSC amplitude and frequency in Tpm4.2^-/-^ neurons (Figure 4A-J) suggest a role for Tpm4.2 in maintaining synaptic strength. The decreased strength of individual synapses may attenuate the depolarization induced by network activity and number of APs generated by a network event. The observed impairment in GluA1 receptor recycling (Figure 2C) is consistent with this idea. It is feasible to suggest that Tpm4.2 depletion also affects the recycling of other receptors, including ion channels which could account for the observed differences in passive membrane properties and mEPSC kinetics.

Despite observing an altered synaptic activity in Tpm4.2^-/-^ primary hippocampal neurons, we did not observe any significant differences in basal synaptic function and plasticity of Tpm4.2^-/-^ brain slice cultures. One possible factor underlying this discrepancy is that slice culture experiments are performed in animals that are 6-8 weeks old, while dissociated cultures are harvested from embryonic mouse brains, and experiments performed at 20DIV. It is possible that Tpm4.2 is only critical for synaptic activity during development, or that adult brains develop other compensatory mechanisms to account for the loss of Tpm4.2. Differences in dissociated neuron culture and brain acute brain slice experiments could also be attributed to differences in model complexity, with acute brain slices also containing other cells which would also be depleted of Tpm4.2. Tripartite synapses for example contain neurons and astrocytes, and as astrocytes exchange information with the synaptic neuronal elements and respond to synaptic activity, they could potentially compensate for the loss of neuronal synaptic activity in Tpm4.2^-/-^ neurons and mask this neuronal phenotype. Future studies could explore this through generating mice with neuron-specific Tpm4.2 knock-out.

An additional contributor to neuronal signaling probability and strength, is the morphology of the neuron and its neurites. Tpm4.2-depleted neurons demonstrated increased total axonal length, increased dendritic complexity early in development, with increases in the number of primary branches per tree, dendrite length and number of interactions near the soma. The observed increased dendrite length, branching and complexity in Tpm4-/- neurons could strongly influence synapse dispersal along a dendrite [22], which in turn affect post-synaptic strength and excitability of neurons. On the other hand, with neurite arborization strongly influenced by extracellular signals and synaptic activity [23], increased dendrite branching and complexity may be a result of hyperexcitable Tpm4.2^-/-^ neurons with increased firing frequency. Further, these increases in dendrite complexity could contribute to the increased network connectivity displayed in live neuron imaging of calcium in Tpm4.2^-/-^ neurons.

On the molecular level, the effect of Tpm4.2 depletion on neurite growth may be associated with its property of cooperatively binding with tropomodulin (Tmod) to filamentous actin. Tropomyosins binding enhances tropomodulin’s capping activity of actin filament pointed ends [24], assisting in the stabilization of filaments and inhibit their disassembly and turnover [25]. Tropomodulins also have isoform specific effects, with Tmod3 not demonstrated to affect neurite outgrowth, while Tmod1 and Tmod2 are reported to increase dendritic complexity [26], and both Tmod1 and Tmod2 require both of their actin-binding sites to regulate dendritic morphology and dendritic spine shape [27]. Tpm4.2 may interact with Tmod1 or Tmod2 and therefore Tpm4.2 depletion may affect this interaction, and result in the observed increase in dendrite branching complexity of Tpm4.2^-/-^ neurons [28]. Neurite outgrowth is therefore proposed to be tightly regulated by numerous components of the actin cytoskeleton, with an isoform-specific function of Tpm4.2 in dendritic outgrowth and complexity.

Dendritic spine expression and morphology can also strongly influence neuron excitability and signaling strength. Dendritic spines are highly dynamic structures that can grow, change shape, and be eliminated through neuronal development, maturation and maintenance. Understanding their morphological properties in different neuron populations can provide insight into their functional differences, with their activity-dependent plasticity directly affecting neuronal signaling, learning and memory processes. Tpm4.2 knock-out neurons exhibit longer, thinner dendritic spines when compared with Tpm4.2^+/+^ neurons. Thin spines are reported to form more transient and weaker synapses [29], and are associated with stress [30], as are longer spines [31]. These longer thinner spines may account for the weakened synaptic activity observed in the dissociated primary hippocampal Tpm4.2^-/-^ neurons. Long thin spines are immature spines which then transform into mushroom or stubby spines following LTP or LTD respectively [32,33]. Here, we show that subtle alterations in normal, healthy dendritic spine dynamics of single cells can have substantial effects on neuronal signaling and learning and associative memory processes such as those we observe in Tpm4.2^-/-^ neurons. These neuronal changes, however, may be masked or mitigated by other processes in more complex brain tissues, including the interaction of glial cells and, in particular the role of astrocytes in regulating neuronal activity and plasticity.

Neuronal hyperexcitability was observed in Tpm4.2^-/-^ neurons, which likely arose due to the increased dendritic complexity and alterations in excitatory surface receptors. These changes, in addition to alterations in spine morphology can result in more aberrant neuronal connections within the hippocampus and neighboring regions [34], depicted in the current study with increased neuronal connectivity in Tpm4.2^-/-^ neurons. Hyperexcitable neurons, and increased neuronal connections are both implicated in producing anxiety-like behaviors [35]. Here, we observed a heightened anxiety phenotype demonstrated in Tpm4.2^-/-^ mice, particularly in female mice. As increasing evidence is implicating hippocampal hyperexcitability with anxiety like behaviors [35], it is important to further investigate Tpm4.2 expression and regulation in the context of models of anxiety could uncover some potential mechanisms underlying anxiety-like behaviors. As the Tpm4.2 was deleted throughout the entire brain in our model, it is important to consider the involvement of other brain regions in the production of the observed anxiety-like phenotype, specifically other limbic regions such as the amygdala and nucleus accumbens [36].

Tpm4.2^-/-^ mice also displayed impaired social memory or socialization with reduced nosing on the social preference test. Impaired recognition memory was also apparent in Tpm4.2^-/-^ mice, with no preference in the Novel object recognition testing, however only females appeared to be affected. These behavioral phenotypes predominantly affecting females, suggests further studies should more closely investigate sex differences in not only Tpm4.2, but other actin cytoskeleton proteins and their functions.

## Conclusion

In conclusion, our study sheds light on the role of Tpm4 neuronal development, signaling and maintenance, highlighting its important regulation in the healthy brain. The observations regarding heightened neuronal firing and neuronal networks producing anxiety-like phenotypes could implicate the dysregulation of Tpm4.2 in anxiety disorders. Future research is required to further dissect the molecular interactions of Tpm4.2 that drive the observed functional changes in early developing and mature neurons. Further, the observation of impaired social and recognition memory warrants further investigation of a role for Tpm4.2 in memory disorders such as dementia.

## 4. Materials and Methods

### 4.1 Generation of Tpm4.2^-/-^ mice

Tpm4.2^-/-^mice were designed as a homozygous knock-out of exon 1b of Tpm4.2 using the C57/Bl6 mouse strain as described in [6]. Mice were genotype*d* by using isopropanol-precipitated DNA from tail biopsies as a template for polymerase chain reaction (PCR), using the following primers Tpm4.2 knock-out Forward primer: 5’- GTGACCTCATGGGCCTGAC-3’ and Tpm4.2 knock-out reverse primer: 5’-GGACGAAAAGTGGGATCG-3’. All procedures involving animals were approved by the UNSW and Macquarie University Animal Care and Ethics Committees and conducted in accordance with national and international guidelines.

### 4.2 Characterization of Tpm4.2^-/-^mice

#### 4.2.1 Perfusion and tissue collection

To confirm the complete knock-out of Tpm4.2, Tpm4.2^-/-^ and Tpm4.2^+/+^ control mice were euthanized and transcardially perfused with 1x phosphate buffered saline (PBS) and their brains harvested. Brains were sagittal sectioned into two halves along the midline and one half was immediately snap frozen in liquid nitrogen and later stored at -80°C for downstream molecular studies.

#### 4.2.2 Western Blot

Snap frozen half brain tissues were lysed with ice-cold Radioimmunoprecipitation Assay (RIPA) lysis and extraction buffer (20mM Tris pH 8.0, 1% Nonidet P-40, 0.25% Sodium deoxycholate, 0.1% Sodium dodecyl sulphate, 150 mM Sodium chloride, 5 mM Ethylenediaminetetraacetic acid, 30mM Sodium fluoride, 60 mM β-Glycerophosphate, 20 mM Sodium pyrophosphate, 1 mM Sodium orthovanadate, 1 mM Dithiothreitol, 1x protease inhibitor, and Milli-Q water). In brief, tissues were thawed upon addition of RIPA buffer followed by sonication at 1pulse/sec with 20% amp in a probe sonicator for 30 secs and then centrifuged at 14,000 rpm for 10 mins at 4 °C. The supernatants were collected and bicinchoninic acid assay was performed for protein quantification using Pierce BCA Protein Assay Kit (Thermo Scientific, Cat No. 23225). 12.5 μg or 10 μg of total protein from brain tissue lysates were loaded onto 12.5% SDS-polyacrylamide gels for brain-regions specific expression and Tpm isoform compensation analysis, respectively. Electrophoresis was run at 80 V for 20 mins through the 4% stacking gel, before 120 V for 2.5 hrs through the 12.5% resolving gel. Resolved proteins were then transferred on to methanol activated polyvinylidine difluoride membrane (Merck Millpore) using a Bio-Rad Trans-blot Turbo transfer system and chilled transfer buffer containing 10% Tris-Glycine SDS buffer (10x), 20% methanol in milli-Q water. Membranes were then blocked with 5% skim milk in Tris buffered saline with 0.1% Tween 20 (TBST) on the rocker for 2 hrs at room temperature followed by incubation with primary antibodies diluted in blocking buffer [Rabbit polyclonal anti-tropomyosin 4.2 (Gift from Peter Gunning, 1:2000) for 3 hrs in 2.5% blocking buffer at room temperature; mouse anti-Tpm3.1/2 (2G10.2, 1:250 Gift from Peter Gunning), mouse anti-total Tpm3 (CG3, 1:250,) and mouse anti-Tpm1.10/12 (clone 5-54, 1:250) in 5% blocking buffer and overnight incubation at 4 °C on rocker) [37]. After 4 washes with TBST (5 mins each at room temperature) the blots were incubated with the respective secondary antibodies (1:5000 in their respective blocking buffer) for 1.5 hrs at room temperature on rocker before washing again 4 times with TBST (5mins each). Next, the blots were developed with Crecendo substrate (Millipore) and visualized using ChemiDoc imaging system (BioRad). All blots were subsequently probed with Glyceraldehyde 3-phosphate dehydrogenase (GAPDH) housekeeping gene (Merck-Millipore; Cat No. MAB374;) (1:5,000 for Tpm4.2 blot and 1:1,000 for all other blots) and imaged to obtain the corresponding loading controls. The western blots were quantified using ImageJ software (NIH).

All values are expressed as mean ± SEM (standard error of mean) and were subjected to unpaired t-test using Graph Pad Prism (Version 8.3). Significance (if found) was indicated as p < 0.05.

### 4.3 Primary Culture of Mouse Hippocampal Neurons

Mouse primary hippocampal neurons were cultured from embryonal day 16.5 (E16.5) from Tpm4.2^+/+^ and Tpm4.2^-/-^ mice with C57/Bl6 background as previously described [38]. In brief, brains were removed from E16.5 mouse embryos and placed in Hanks buffered salt solution (HBSS; Sigma). Meninges were removed and hippocampi extracted using microclippers. A 1:100 dilution of Trypsin (Sigma-Aldrich) was then added to hippocampi and incubated at 37 °C for 20 mins. Deoxyribonuclease I 1:100 final concentration of 0.1 mg/mL; Sigma) was added for 30 secs before washing twice with 10 mL of DMEM (Life Technologies, Berkley, CA, USA) with 10% fetal bovine serum (FBS, Thermofisher Scientific) to remove DNAseI. Hippocampi were then dissociated by trituration with 1 mL DMEM/10% FBS using fire-polished, serum coated Pasteur pipettes. Cells for Receptor recycling assay and calcium imaging of live neurons were then plated on poly-D-lysine (PDL; Sigma)-coated 1.5 mm glass coverslips at a density of 70,000 cells per well in a 24-well plate and incubated at 37 °C and 5% CO_2_ for 2 hrs in DMEM/10% FBS. Media was then changed to Neurobasal medium (NBM; Neurobasal, Life Technologies; supplemented with 2% B27, Life technologies and 0.25% GlutaMAX, Invitrogen).

### 4.4 Receptor internalization assay

#### 4.4.1 Neuron treatment

Primary hippocampal neurons were seeded at 70,000 per well on 1.5 mm, PDL-coated glass coverslips and incubated in NBM at 37 °C with 5% CO_2_. For 18 days. a receptor internalization assay was performed as previously described [39]. In brief, 80% of culture media was removed and neurons either left untreated, incubated with Extracellular Solution (ECS; 150 mM NaCl, 2 mM CaCl_2_, 5mM KCL, 10 mM HEPES (pH7.4) and 30 mM glucose in dH_2_O), 25 μM NMDA in ECS, 100 M Glycine in ECS or 25 M Bicuculline in ECS for30 mins at 37 °C with 5% CO_2_. Incubation solutions were then aspirated and replaced with original cell culture medium and reincubated for 60 mins before fixing for 5 mins in 4% PFA.

#### 4.4.2 Surface and total GluA1 receptor probing

Half of the neurons were then probed for surface GluA1 receptors only by blocking for 1 hr at RT in blocking buffer (2% FBS (Sigma) in PBS) followed by incubation for 60mins with mouse N-terminal anti-GluA1 (1:250 in BB; Merck-Millipore MAB2263) diluted in blocking buffer. Surface probed neurons are then washed in PBS and fixed with 4% PFA for 5 mins, permeabilized with 0.1% Triton-X in PBS for 5mins at RT with PBS wash steps in between each. The remaining half of the neurons are probed for total GluA1 receptors by permeabilized with 0.1% Triton-X in PBS for 5 mins at RT, washing in PBS, blocking for 60 mins in BB before incubation with:150 mouse anti-GluA1 at RT for 60 mins. All neurons are then washed, blocked for 60 mins in blocking solution and incubated with Chicken anti-MAP2 (1:500 in blocking buffer; Abcam ab5392) for 4 °C overnight. All neurons are then washed and incubated for 60 mins with donkey anti-mouse AlexaFlour-555 (1:500 Life Technologies in BB) for GluA1, goat anti-chicken AlexaFlour-647 (1:500; Life Technologies in BB) for MAP2. Secondary antibodies are then washed off in PBS before incubation with phalloidin 488 (1:100 in PBS, Thermo Fisher Scientific) and DAPI (1:1,000, Life Technologies) for 20 mins a RT. Neurons are then washed in PBS, dipped in H_2_O and mounted onto glass slides with Fluromount G (Thermo Fisher Scientific)

#### 4.4.3 Imaging and analysis receptor internalization

Slides were imaged on a Zeiss AxioImager, using a 63x oil objective. Neurons were stitched, exported as uncompressed .Tif files and analyzed on FIJI (Image J). Relative intensity was measured for 3 dendrites per neuron and 10 neurons per condition, maintaining a consistent distance away from the soma (5 M) and measurement area (15uM^2) and normalized to background intensity. Ratios of surface to total GluA1 intensity were measured and the ratio compared between groups using GraphPad Prism software (version 10.2)

### 4.5 Plasmid production and cloning

For calcium imaging of live neurons; we used a the green fluorescent protein (GFP) based GCaMP sensor protein with fast kinetics jGCaMP7f as previously characterized [40] and inserted it into an AAV vector contained a PhP.B capsid, human Synapsin promoter and woodchuck hepatitis virus post-transcriptional regulatory element (WPRE), flanked by AAV inverted terminal repeats (ITRs). NEBuilder Hifi DNA Assembly Master mix was used to clone plasmids (E2621L, New England Biolabs, Massachusetts, USA). DNA fragments were amplified by PCR and 0.06 pmol was combined with 0.06 pmol of digested vector fragments, 5 μL of water and 5 μL of NEBuilder DNA Assembly Master Mix. The mixture was incubated for 1 hr at 50 °C before 5 μL was added into 50 μL of One Shot Stbl3 Chemically Competent *E. coli* (C737303, Thermo Fisher Scientific, Massachusetts, USA) and incubated on ice for 30 mins, heat-shocked at 42 °C for 45 secs, then placed back on ice for a further 2 min. Transformed bacteria were then mixed with 200 μL of SOC Outgrowth Medium (B9035, New England Biolabs, Massachusetts, USA), incubated with 300 rpm shaking for 1 hr at 37 °C and plated on agar plates with 1:1,000 Ampicillin. Plates were then incubated at 37 °C overnight.

For each construct, a pipette tip was used to pick five distinct colonies from each agar plate and added to LB broth (0.5% Yeast [Sigma Aldrich, Missouri, USA], 1% Tryptone [G-Biosciences, New Delhi, India] and 1% NaCl [Sigma-Aldrich, Missouri, USA]). DNA was then extracted from bacterial culture using Wizard Plus SV Miniprep Purification System (Promega, Wisconsin, USA), as per manufacturer’s protocol. In brief, the culture was pelleted for 5 mins, before resuspension in 250 μL of Cell Resuspension Solution. 10 μL of Alkaline Protease Solution was then added and the mixture incubated for 5 min at room temperature before neutralization with 350 μL of Neutralization solution. The mixture was centrifuged (10 mins at room temperature, 21,000 g) and the supernatant transferred to the Spin Column placed in a Collection Tube, and centrifuged (21,000 g for 1 min at room temperature). Flowthrough was discarded, and the bound DNA was washed with 750 μL of Wash Solution spun (centrifuged for 1 min at room temperature, 21,000g). The wash step was repeated with 250 μL of Wash Solution. The Spin Column was transferred to a sterile 1.5mL microcentrifuge tube and the DNA was eluted by incubating the Spin Column membrane with 50 μL of Nuclease-Free Water for 2 mins at room temperature, followed by centrifuging for 1 min at room temperature.

PureLink HiPure Plasmid Maxiprep Kit (Thermo Fisher Scientific, Massachusetts, USA) was used to obtain larger DNA amounts. Bacteria were grown in 4 mL of LB Broth (1:1,000 ampicillin) for 8 hrs, before expansion to 250 mL LB Broth (1:1,000 ampicillin) overnight. Overnight culture for spun at room temperature for 10 mins at 4,000 g and the pellet resuspended in 10 mL of Resuspension Buffer with RNase A. 10 mL of Lysis Buffer was then added to the cells, mixed by inversion and incubated for 5 mins at room temperature. 10mL of Precipitation Buffer was then added, and the mixture centrifuged at 12,000 g for 30 mins at room temperature. Supernatant was loaded onto the previously equilibrated column and drained by gravity flow. The column was washed with 60 mL of Wash Buffer and drained by gravity flow. 15 mL Elution buffer was then added to the column to elute DNA using gravity flow. The DNA was mixed well with 10.5 mL of isopropanol, and centrifuged at 12,000 g for 30 mins at 4 °C. Supernatant was removed, and pellet resuspended in 5 mL of 70% ethanol, followed by centrifugation at 12,000 g for 10 mins at 4 °C. Supernatant was discarded, and the pellet was left to air-dry for 10 mins. The DNA was resuspended in 300 μL of Nuclease-Free Water and used for adeno-associated virus production.

### 4.6 Production of adeno-associated virus for neuron transduction

GCamp7f-eGFP, Tpm4.2-mRuby2 and Tpm4.2-IRES-mRuby2 were packaged into PHP.B capsids and adeno-associated viruses were produces as previously described [39]. In brief, HEK293T cells were grown to 70-80% confluency in DMEM/10% FBS before replacement with Iscove modified Dulbecco medium (Sigma) + 5% FBS 3 hrs prior to transfection with polyethyleneimine-Max (PEI-max, Polysciences) containing pF0delta06 and AAV-PHP.B plasmid with rep and cap sequences.

### 4.7 Calcium imaging of live neurons

Primary hippocampal neurons for calcium imaging of live neurons were seeded at 70,000 on 1.5 mm, PDL-coated glass coverslips. Neurons were transduced with 1 x 10^12vg/coverslip of AAV-PHP.B jGCamp7f and incubated at 37 °C with 5% CO_2_ for 20 DIV. At each timepoint, cells were imaged at 37 °C with 5% CO_2_ on an Axio Observer 7 Live cell imager (Carl Zeiss, Germany) at 5x magnification for 5 mins with 500 ms intervals. Following live-cell imaging, 5 M ionomycin was added to each well and imaged after 10 mins to obtain baseline fluorescence. Cells were then fixed with 4% paraformaldehyde (PFA) and stored in PBS for immunocytochemistry.

Time-series images were converted to .tif stacks in Image J. Baseline fluorescence images obtained following Ionomyocin incubation were subtracted from the timeseries stack. A somatic region of interest (ROI) was drawn for each active neuron in FIJI [41] and the fluorescent time series for each ROI within in the field of view (FOV) was saved as a table. Time series were analyzed using custom MATLAB (version R2024b, MathWorks) routines adapted from previously published *in vitro* calcium image analysis pipelines [42,43]. Normalized fluorescence intensity (D_F/F_) was extracted using the getDFF function in CaPTure [43] with tau [2 50]. Peaks and their locations were identified using the MATLAB findpeaks function (“MinPeakHeight” = 0.01 D_F/F_,“MinPeakDistance”=4 timepoints). To quantify the synchronous activity the normalized fluorescence intensity for each ROI within a FOV was averaged and the peaks of the average intensity timeseries were extracted as described above.

### 4.8 Electrophysiology

#### 4.8.1 Whole cell patch clamping of primary neurons in dissociated cultures

Primary hippocampal neurons were seeded at 70,000/well in a 24-well plate onto 1.5 mm PDL coated coverslips and incubated in NBM at 37 °C and 5% CO_2_. At 17 and 18 DIV, Coverslips were transferred to recording bath on a Leica DM IL inverted microscope, which was perfused with extracellular solution (110 mM NaCl, 10 mM HEPES, 10 mM glucose, 2 mM CaCl_2_, 0.8 mM MgCl2, 5 mM KCl) at room temperature using a peristaltic pump (Minipuls 3, Gilson, France). After a cell was patched, the perfusion was switched to a separate 20 ml aliquot 0.5 μM TTX and 100 μM picrotoxin. Patch electrodes were made from glass capillaries 1.2 mm OD, 0.94 ID, 100 length (Harvard Apparatus), pulled using Narishige Model PC-10 microelectrode puller to a tip resistance of 3-5 MΩ. Electrodes were filled with internal solution (110 mM cesium methane sulfonate, 8 mM NaCl, 10 mM HEPES, 2 mM Mg_2_ATP, 0.3 mM Na_3_GTP, 0.1 mM spermine tetrahydrochloride, 7 mM phosphocreatine, 10 mM EGTA, 5 mM CsCl) with 50 μM Alexa Fluor 594, filtered through a 0.22 μm syringe driven filter unit (Millex). Recordings of mEPSCs were made at a holding potential of -70 mV with an Axopatch 200B amplifier, filtered at 2 kHz, digitized at 5 kHz with a Digidata 1440 A, and saved with Clampex 10.2 (Molecular Devices, USA).

#### 4.8.2 mEPSC analysis in primary neurons of dissociated cultures

mEPSCs were detected and measured using Axograph (Sydney, Australia). A notch filter (49.9-50.1 Hz) was applied and an event template (maximum 0.5 ms rise time, 3 ms minimum decay time) was used to detect mEPSC[44]. Events outside of 5-150 pA were excluded, and detected events were manually verified. Event amplitude and inter-event interval was measured. Either the first 1,000 events or 5 mins of activity were analyzed from each cell, as there was a large variability in activity frequency.

#### 4.8.3 Changes of field potential recording in brain slices

All animal studies were carried out in accordance with the New South Wales Animal Research Act and Regulation and approved by the Animal Ethics Committee of UNSW Sydney under ethics protocol numbers 14/113A and 18/37B. Mice were housed in a temperature-controlled facility (22-24 °C) on a 12-hour light-dark cycle. For all experiments, mice of both sexes were used at 6-8 weeks old. Data collection was performed blind to genotype, interspersed so no more than two animals of the same genotype were used on consecutive days.

Mice were anesthetized by open-drop exposure to isoflurane in an induction chamber before decapitation and the brain removed and placed in ice-cold modified artificial cerebrospinal fluid (ACSF; 124 mM sucrose, 62.6 mM NaCl, 2.5 mM KCl, 26 mM NaHCO3, 1.2 mM NaH_2_PO_4_, 10 mM glucose, 0.5 mM CaCl2, and 3.3 mM MgCl_2_). The brain was hemisected and immersed in cold modified ACSF. Horizontal slices (400 μm) were cut using a vibratome (model VT1200, Leica, Wetzlar, Germany) at room temperature. The solution containing the brain slices was continuously infused with 95% O_2_/5% CO_2_. Slices were left to recover for at least 1 hr before recording. Slices were used within 7 hrs of cutting, corresponding to the optimal period of slice health [45].

Slices were transferred individually to the tissue recording system (Kerr Scientific Instruments, Christchurch, New Zealand) and continuously perfused with standard ACSF at room temperature. A bipolar, Teflon-coated tungsten stimulating electrode (Kerr Scientific Instruments) was placed in the stratum radiatum, aligned to the end of the dentate gyrus, and the recording electrode was placed approximately 800 μm from the stimulating electrode. Stimuli were delivered via an isolated stimulator (model DS2, Digitimer, Hertfordshire, England, or A-M Systems, Model 2200, Washington, USA), triggered through a Powerlab (model 4/2ST, AD Instruments, Sydney, Australia) or multifunctional data acquisition card (National Instruments NI PCI-6221, USA or Data Translation DT9816; USA). Field potentials were amplified at 100x using a KSI Tissue Recording System Amplifier (Kerr Scientific Instruments, Christchurch, New Zealand) and digitized with the Powerlab or multifunctional data acquisition card. Traces were acquired using Scope (AD Instruments, Sydney, Australia), AxoGraph X (Axograph Scientific) or custom software. To determine the optimal electrode positions, stimuli (15-20 V 100 μs) were delivered every 8-10 secs and electrodes were lowered until an extracellular field excitatory synaptic potential (fEPSP) was observed. The depth of the recording and stimulating electrodes were then adjusted to produce the largest response. After finding the optimal electrode position, the stimulus frequency was reduced to once every 30 seconds and left to stabilize for 10 minutes. A stimulus response curve was then conducted by varying the stimulus intensity from 5 to 70 V. Slices were discarded if the maximum fEPSP amplitude was below 0.6 mV as the small response was considered to be a marker of poor health. The stimulus intensity eliciting 50% of the maximum amplitude was identified, and field potentials were evoked in pairs with a 50 ms interval at this stimulus intensity at 30 s intervals. After obtaining a stable baseline for 20-30 minutes, LTP was induced with 2 bursts of high frequency stimulation (100 Hz 1s). For LTD experiments, LTD was induced with 900 paired pulses (50 ms interval) at 1 Hz. Responses were then recorded for the following 60 minutes.

Electrophysiological data were analyzed offline using AxoGraph (Sydney, Australia). Measures included the fibre volley amplitude, the fEPSP slope and the paired pulse ratio. Fiber volley amplitude was defined as the amplitude of the negative peak preceding the fEPSP. The fiber volley amplitude is indicative of the number of presynaptic axons activated by stimulation. The fEPSP slope was defined as the maximum slope during the initial 2.5 ms after the fiber volley. This was used to measure excitatory activity. Slope was measured instead of amplitude, as the amplitude can be contaminated by the population spike at higher stimulus amplitudes. The paired pulse ratio was calculated as the second fEPSP slope divided by the first, to give a measure of presynaptic transmitter release probability [46].

#### 4.8.4 Statistical analysis for electrophysiology

Statistical comparisons were performed using GraphPad Prism (GraphPad Software, La Jolla, USA). The D’Agostino & Pearson test was used to test data sets for normality. Differences between groups were tested using repeated measures ANOVA, paired t-test, unpaired t-test, Mann-Whitney test, or Kolmogorov-Smirnov test as indicated with α = 0.05. Data are presented as mean ± standard error of the mean. Data collection and analysis were performed blind to the genotype of the animals.

### 4.9 Neurite outgrowth

#### 4.9.1 Neuron plating and transfection for neurite outgrowth analysis

For neurite growth experiments, primary hippocampal Tpm4.2^+/+^ and Tpm4.2^-/-^ neurons were seeded at 70,000 cells/well in a 24-well plate containing PDL-coated 12mm glass coverslips and cultured in 1 ml per well in complete NBM at 35 °C and 5% CO_2_.

For neurite tracing experiments, pEGFP-C1 plasmic (Clontech) was used for control neurons. Tpm4.2^+/+^ and Tpm4.2^-/-^ neurons were transfected with pEGFP-C1 at the 2 DIV with Lipofectamine 3000 (Thermo Fisher Scientific, Massachusetts, USA) by making a transfection mixture containing 0.5 µg of DNA and 1 µL of Lipofectamine 3000 in a total of 100 µL NBM per well of a 24-well plate. The transfection mixture was incubated with the cells for 90 mins at 37 °C and 5% CO_2_ in the 50% of the initial culture media volume (while the other 50% was collected before transfection procedure and kept at 37 °C). After incubation, the media was aspirated, and conditioned media was added to the cells.

#### 4.9.2 Immunocytochemistry for neurite outgrowth analysis

Two days after transfection at 4 DIV, neurons were fixed with 4% paraformaldehyde (PFA) at room temperature for 15mins, washed in PBS and permeabilized in 0.1% Triton-X (Sigma-Aldrich) for 5 mins at room temperature. Neurons were then washed and blocked in BB (2% FBS in PBS) at room temperature for 1 hr before incubation in the following primary antibodies diluted in BB - mouse anti-Tau1 (1:500 Millipore ab3420), chicken anti-B3-tubulin (1:250, Millipore ab9354) overnight at 4 degrees. The following day, primary antibody was washed off in PBS and secondary antibodies donkey anti-mouse Alexa-555 and goat anti-chicken Alexa-647 (both 1:500, Life technologies) were diluted in BB and placed on neurons for 1hr at room temperature. Neurons were washed in PBS and mounted on glass microscope slides with Prolong Gold antifade reagent with DAPI (Life technologies).

#### 4.9.3 Imaging of neurite outgrowth

Tpm4.2^+/+^ and Tpm4.2^-/-^ mouse primary hippocampal neurons transfected with pEGFP-C1 were imaged, using an Axio Imager upright fluorescent microscope fitted with a monochrome camera, with EC-Planachromatic Neofluar 40x magnification objective (NA 0.75, WD 0.71mm, Air immersion). Fluorescent illumination was obtained with a Xenon HXP lamp. The fluorescent filter sets used were BP450-490/BS495/BP500-550 (FS#38) for 488 nm channel, BP533-558/BS570/BP570-640 (FS#43) for 555 nm channel, and BP625-655/BS660/BP665-715 (FS#50) for 647 nm channel. At least 20 transfected neurons were imaged per single coverslip, and four coverslips per single experimental group were imaged per one biological replicate (80 imaged neurons per biological replicate in total – 3 biological replicates). Images are representative of all three biological replicates.

#### 4.9.4 Morphological and statistical analysis for dendritic spines

The axonal compartment of Tpm4.2^+/+^ and Tpm4.2^-/-^ neurons, transfected with pEGFP-C1 plasmid was identified as tau1-positive/β3-tubulin-positive. The dendritic compartment was verified as tau1-negative/β3-tubulin-positive. For each experimental group, ∼20 neurons (∼5 neurons per coverslip) from 80 imaged neurons per biological replicate were chosen, using Random number generator (RNG). The images were processed in ImageJ (v.2.1.0). The morphological analysis of neurons was performed, using the semiautomated approach in Neurolucida software (MBF Bioscience, Vermont, USA, v2019.1.1) to outline soma, axons, and dendrites. AutoNeuron workflow was used to initiate automatic tracing, followed by manual corrections, labelling, and branch ordering. To quantify traced axonal and dendritic compartments in Neurolucida, Branched Structure Analysis and Centrifugal Sholl Analysis options in Neurolucida Explorer package were used. Statistical analysis was performed in GraphPad prism software (version 9.1.2). To determine if experimental groups have normal Gaussian distribution, we performed Anderson-Darling, D’Agostino-Pearson, Shapiro-Wilk, and Kolgomorov-Smirnov normality tests. After these tests have shown no Gaussian distribution within the experimental groups, the significance was determined with non-parametric Mann-Whitney U test for Tpm4.2^+/+^ and Tpm4.2^-/-^ neurons transfected with pEGFP-C1.

#### 4.9.5 DiI injection for dendritic spine analysis

Cells were fixed in 4% PFA for 15 mins at room temperature and washed with PBS. The fixed coverslip was placed in a bath of PBS on a Leica DMIL microscope. A 1% solution of 1,1′-Dioctadecyl-3,3,3′,3′-tetramethylin-docarbocyanine perchlorate (DiI) in ethanol was loaded into a sharp microelectrode. The tip of the microelectrode was maneuvered into the target cell soma and left for five minutes to allow dye to enter the cell. Sometimes pressure was added to expel the DiI solution. Cells were randomly selected for injection.

Approximately ten cells per coverslip were injected. The coverslips were then left in 1x PBS at 4 °C overnight, or at room temperature for five hours to allow the DiI to diffuse into neuronal processes. The coverslips were then mounted with either Fluoromount or Prolong Gold, then sealed with nail polish the next morning before imaging.

#### 4.9.6 Imaging of dendritic spine analysis

Immunofluorescence images were taken using an LSM 710 confocal microscope (Zeiss) with a 63x 1.4 NA oil immersion objective. DiI was excited with a 561 nm DPSS laser and emission captured at 519-673 nm. For dendritic spine morphology analysis, two 20 μm length segments of secondary dendrites, 50-75 μm away from the cell soma were selected from each transfected neuron. A Z-stack image with 0.21 μm interval and 0.04 x 0.04 μm pixel size was acquired of each selected dendrite.

Deconvolution was performed on images using the DeconvolutionLab2 plugin in ImageJ [47]. Images were processed with 5 iterations of the Richardson-Lucy algorithm using a point spread function generated by Diffraction PSF 3D plugin (Optinav).

#### 4.9.7 Spine quantification

To investigate whether Tpm4.2 deletion affects spine morphology, primary neuronal cultures were prepared from Tpm4.2^+/+^ and Tpm4.2^-/-^ mice, fixed at 17-18 DIV, stained with *Dil* to assist labelling of entire cell membranes, and imaged. 51 cells were analyzed for the Tpm4.2^+/+^ group, and 57 cells were analyzed for the Tpm4.2^-/-^ group, from 3 separate culture preparations.

Spine morphology was analyzed according to Risher et al., 2014 [48]. Z-stack images were imported to RECONSTRUCT [49]. The length of the dendrite, and the length and widths of spines were manually traced and measured. Measurements were copied to an Excel spreadsheet to calculate the spine density, average protrusion width, and average protrusion length for each cell that was imaged. Data were analyzed and graphs were generated in GraphPad Prism. Data are given as mean ± SEM, unless otherwise indicated.

### 4.10 Mouse housing for Behavioral analysis

Male and female Tpm4.2^-/-^ (n = 20) and Tpm^+/+^ mice (n = 25) littermates with C57BL/6 background were transported at least two weeks prior to testing for habituation from Macquarie University and housed on arrival in the animal facility at the School of Medicine, Western Sydney University (Campbelltown campus, Australia). Adult A/JArc mice were used in the social preference test to trigger explorative behavior of test mice. All animals were group housed (2–3/cage) in individually ventilated cages (Type Mouse Version 1; Airlaw, Smithfield, Australia; air change: 90–120 times per hour averaged; passive exhaust ventilation system), containing corn cob bedding, a mouse igloo (Bioserv, Frenchtown, USA), and a crinkle nest to provide nest building opportunities (Crink-l’Nest, Tecniplast, Australia). Mice were kept under a 12:12 h (light phase: 0900-2100 h with white light at an illumination of 124 lx; dark phase: 2100-0900 h with red light at an illumination of less than 2 lx), food and water ad libitum, with a temperature between 22-24°C and humidity between 40-60 RH. Mice were 7 months old at the start of behavioral experiments. Research and animal care procedures were approved by the Western Sydney University Animal Care and Ethics Committee (approval number: A12918) and were in accordance with the Australian Code of Practice for the Care and Use of Animals for Scientific Purposes.

### 4.11 Behavioral Phenotyping

For habituation purposes, all test animals were transported to the testing room 30 mins prior to behavioral testing and all experiments were performed within the first 6 hrs of the light phase, with an inter-test interval of at least 48 hrs. All test equipment was cleaned after each trial with 80% ethanol solution. Tests were carried out in the following order: Open field, novel object recognition, social preference test, elevated plus maze, prepulse inhibition and fear conditioning. The sample size for each experimental test condition was n = 10-12.

#### 4.11.1 Open field (OF)

Locomotor activity, exploration and anxiety-like behaviors were measured in the OF and were conducted as previously published from our laboratory [50,51]. Mice were placed into the right corner of the OF chamber (43 x 43 cm; Activity Monitor, Med Associates Inc., Fairfax, USA) and allowed to explore the arena freely for 30 mins. The arena was divided into a central and peripheral zone (MED software coordinates 3/3, 3/13, 13/3/, 13/13) with the central zone being a more aversive, anxiety-inducing zone of the open field [52]. Software settings for the detection of locomotion were box size: 3; ambulatory trigger: 2; resting delay: 1,000 ms; resolution: 100 ms. Time, horizontal (distance travelled) and vertical activity (rearing) in central and peripheral zones were measured by the chambers’ infrared photo beams. The ratio of central to total distance travelled (distance ratio) and time (time ratio) spent in the central area of the OF were taken as measures of anxiety.

#### 4.11.2 Novel object recognition test (NORT)

The distinction between familiar and unfamiliar objects is an index of recognition memory and is measurement by the innate preference of rodents for novel over familiar objects [53]. The protocol was adapted from previous publication from our group [50]. The apparatus for NORT was a grey perspex arena (35 x 35 x 30 cm) and the protocol ran over two consecutive days. On the first day, mice were habituated twice to the test chamber for 10 mins and allowed to freely explore the empty arena, with an intertrial interval of 2 hrs. On the second day, mice were placed in the NORT arena with two identical objects (2 DUPLO® elephants or 2 blocks of DUPLO®) and allowed to explore the objects for 10 mins. After a 15 min intertrial interval, mice were replaced in the arena with one familiar object and one novel object (LEGO® elephant + block of LEGO®) for 10 mins. Object exploration was scored when mice exhibited nosing behavior towards the objects (i.e. when the mouse directed its snout towards an object at a distance of < 1 cm). Object recognition was reported as INDEX (time spent nosing the novel object expressed as a percentage of the time spent nosing the novel + familiar objects); the behaviour was manually scored using ANY-maze™.

#### 4.11.3 Social preference test (SPT)

The SPT test was used to assess sociability and social novelty preference (i.e. social recognition memory) in test mice and were conducted as previous publication from our group [50]. The apparatus consists of three chambers, i.e. a central chamber (around 9 cm x 18 cm x 20 cm) and two outer chambers (around 16 cm x 18 cm x 20 cm). The dividing walls were made of clear Plexiglas, with square passages, 4 cm high and 4 cm wide. One circular cage (i.e. mouse enclosure) was placed into each outer chamber. The mouse enclosures were 15 cm in height with a diameter of 7 cm and bars spaced 0.5 cm apart to allow nose contact between mice but prevent fighting. The chambers and enclosures were cleaned in-between trials (inter-trial interval of 2 mins) and fresh bedding was added prior to each mouse. During the habituation trial, mice were placed individually in the central chamber and allowed to freely explore the apparatus and the two empty enclosures for 5 min. For the sociability test, an unfamiliar adult age-matched A/J mouse was placed in one of the two enclosures, while the experimental mouse was enclosed in the central chamber. Then the test mouse was allowed to explore all three chambers for 10 mins. Finally, test animals were observed in a 10 mins social recognition test. For this, a second, unfamiliar A/J mouse was placed in the previously empty chamber so that the test mouse had the choice to explore either the familiar A/J mouse (from the previous trial) or the novel, unfamiliar mouse. AnyMazeTM tracking software was used to determine the time spent in the different chambers, number of entries, and distance travelled. In addition, time spent sniffing the opponent (i.e. A/J mouse) was recorded manually (i.e. snout of test mouse within the enclosure containing the opponent mouse or < 1 cm away from enclosure).

#### 4.11.4 Elevated plus maze (EPM)

The EPM represents the natural conflict between the tendency of mice to explore a novel environment and the tendency to avoid a brightly lit open areas [54]. The behavior is also influenced by thigmotaxis and the fear of heights. The EPM was in the shape of a ‘+’, with the four arms extend from a central platform and raised 1 m above the floor. Two alternate arms were dark and enclosed (30.5 cm × 6.5 cm, sidewall height 18.5 cm) while two alternate arms were open (30.5 cm × 6.5 cm, no sidewalls, illumination 40 lx) with a central platform connecting the arms (6 cm × 6 cm). The mouse was placed onto the center field of the EPM (faced to a closed arm) and was allowed to freely explore the maze for 5 min. AnyMazeTM tracking software was used to determine the time spent in open/closed arms, number of entries and distance travelled by the test mice. Anxiety was measured by the time spent on open arms as well as ratio of open arm entries and distance (compared to total number of entries / total distance travelled). These parameters are inversely related to anxiety. The number of total arm entries / total distance travelled reflects general motor activity (locomotion).

#### 4.11.5 Fear-conditioning (FC)

Fear conditioning (FC) is a type of associative learning task in which mice learn to associate a particular neutral conditional stimulus (CS) with an aversive unconditional stimulus (US) and show a conditional response [55]. The FC protocol used here evaluates both cued and contextual conditioning to assess amygdala-dependent and hippocampal-dependent fear-associated memory respectively. On day 1 (training), animals were placed into the test chamber (fear conditioning chambers from MED Associates Inc.) with vanilla scent cue presented in the chamber and lights on. For the first 2 mins, mice were left undisturbed and could explore and habituate to the environment. After 2 mins, the conditioned stimulus (CS: 30 s duration, 80 dB tone stimulus) was paired with a co-terminating unconditioned stimulus (US: electric foot shock of 0.4 mA for 2 s duration) twice with an inter-pairing interval of 120 secs. The test mouse was returned to its home cage 120 s after the second CS–US pairing. On day 2 (context test), the mouse was returned to the testing chamber with the scent cue and light present, but no sound, for a total of 7 mins. On day 3 (cue test), mice were placed in an altered context (e.g. no scent cue present, and a black wall in triangle was placed into the chamber, but lights are still on). Following the first 120 s, during which no auditory stimulus was presented (pre-CS), the CS was then presented continuously for 5 mins. The experiment was then terminated after another 120 secs without CS. In all trials, the percentage freezing response (the absence of all but respiratory movement) per 1-min block was automatically measured using SOF-843 video freeze (MED Associates Inc.), with a freezing threshold of 10. Total averages were calculated for each day, as well as the average freezing per minute in the cue test, during baseline testing in the first 2 mins versus cue freezing in minutes 3 to 7.

#### 4.11.6 Prepulse inhibition (PPI)

Prepulse inhibition (PPI), an operational measure of sensorimotor gating, is the attenuation of the startle response by a non-startling stimulus (prepulse) presented 30 – 500 ms before the startling stimulus (startle pulse). Startle reactivity can be measured using SR-LAB startle chambers (San Diego Instruments, San Diego, USA) where the startle response intensity of rodents (whole body flinch amplitude) can be measured using a piezoelectric accelerometer. As previously described, animals were habituated to the startle chambers and the test enclosures twice a day for 10 min on two consecutive days (with an intertrial interval of 2 hrs). On the third day, the PPI test was carried out and consisted of a 5-min acclimatization period to a 70 dB background noise, followed by 97 trials presented in a pseudorandom order to test the acoustic startle response (ASR) and PPI: 5 × 70 dB trials (background noise); 5 × 100 dB trials; 15 × 120 dB trials (for ASR) and 72 PPI trials comprising six sets of a pre-pulse of either 74, 82 or 86 dB presented 32, 64, 128 or 256 ms (variable interstimulus interval; ISI) prior to a startle pulse of 120 dB. The intertrial interval varied randomly between 10 and 20 secs. Responses to each trial were calculated as the average mean amplitude detected by the accelerometer. The startle response was calculated as the mean amplitude across all 15 startle trials and percentage PPI (% PPI) was calculated as [(mean startle response (120 dB) – PPI response)/mean startle response (120 dB)] x 100. For acoustic startle habituation, blocks of the acoustic response to 120 dB startle pulses presented at the beginning, in the middle and at the end of the PPI protocol (i.e. averaged across 5 trials each) were used to determine the effect of ‘startle block’ [56].

#### 4.11.7 Statistical analysis

Behavioral data were analyzed using GraphPad PRISM 9.4.0. Three-way repeated measures (RM) or two-way analysis of variance (ANOVA) were conducted for the within subjects factors (time x sex x genotype) and the between subjects factors ‘sex’ and ‘genotype’. Paired t-tests were used for analysis between two groups (familiar x new). Single sample t-tests against chance levels (i.e. 50%) were used for the SPT and NORT exploration index. If three- or two-way ANOVA interactions were detected, Bonferroni post-hoc tests were used to follow these up and identify differences between individual groups. Data is presented as mean ± standard error of the mean (SEM), and F-values and degrees of freedom are presented for ANOVAs, Differences were regarded as statistically significant if p < 0.05, trends were mentioned where p = 0.05 - 0.06. Significant “genotype” effects are indicated by “#” (#p < 0.05, ##p < 0.01 and ###p < 0.001), “sex” effects by “+” (+p < 0.05, ++p < 0.01 and +++p < 0.001), “repeated measure” effects by “&” (&&&p < 0.001) and significant interactions by “^” (^p < 0.05, ^^p < 0.01 and ^^^p < 0.001). Significant t-test results against chance level (i.e. 50 %, NORT/SPT) and Bonferroni post-hoc are indicated by “*” (*p < 0.05).

## Supplementary Materials

## Funding

This work was supported by funding from the National Health and Medical Research Council (NHMRC (grant# APP1083209 and grant# APP200660) and the Australian Research Council (ARC) (grant #DP180101473) to TF. RRP is supported by the Ainsworth Medical Research Innovation Fund (AMRIF) as well as Dementia Australia. TK is supported by a Drug Development Grant from FightMND, NSW Health, as well as the AMRIF. SS has been supported by an International Macquarie Research Excellence Scholarship (iMQRES) from Macquarie University.

## Competing Interests

The authors have no relevant financial or non-financial interests to disclose.

## Author contributions

Conceptualization: TF, JP and TK; Methodology: TF, SG, CC, RRP, TT, HS, EP, DY and SS. Formal analysis and investigation: SG, CC, RRP, TT, SS, WG, JP and TF; Writing - original draft preparation: SG, CC and RRP; Writing - review and editing: SG, RRP, CC, JP and TF; Funding acquisition: TF, TK, JP and PG; Resources: TF, PWG, TK, JP and ECH; Supervision: TF, JP and TK. All authors read and approved the final manuscript.

## Data Availability

Additional data will be made available upon request.

## Ethics approval

Approval was obtained from the animal ethics committee of Macquarie University. The procedures used in this study adhere to the tenets of the Declaration of Helsinki.

## Notes

### Competing Interest Statement

The authors have declared no competing interest.

