## Supplementary Figures and Table for "Knock-out of Tpm4.2/actin filaments alters neuronal signaling, neurite outgrowth and behavioral phenotypes in mice": Genoud et al - Supplementary Figures and Table.pdf

Fig. S1

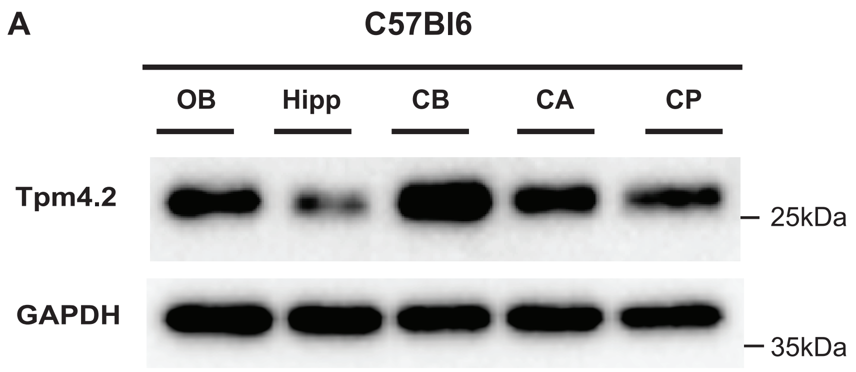

A
C57Bl6
OB
Hipp
CB
CA
CP
Tpm4.2
25kDa
GAPDH
35kDa


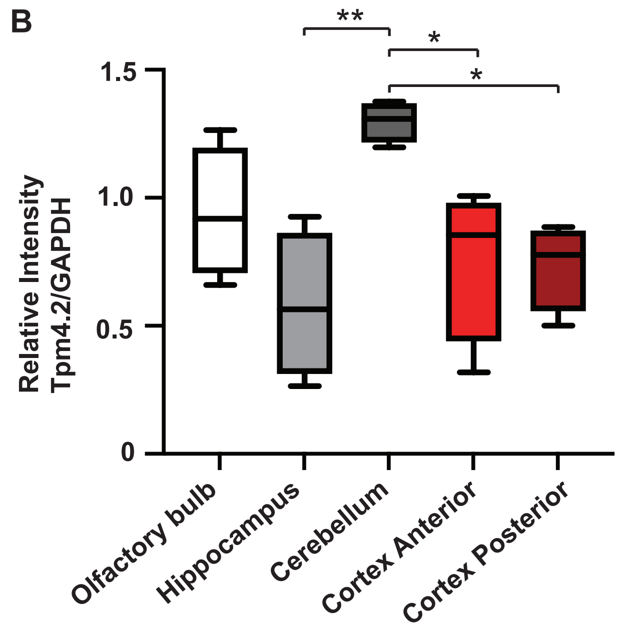

B
**
*
*
1.5
1.0
0.5
0
Relative Intensity
Tpm4.2/GAPDH
Olfactory bulb
Hippocampus
Cerebellum
Cortex Anterior
Cortex Posterior

Fig. S2

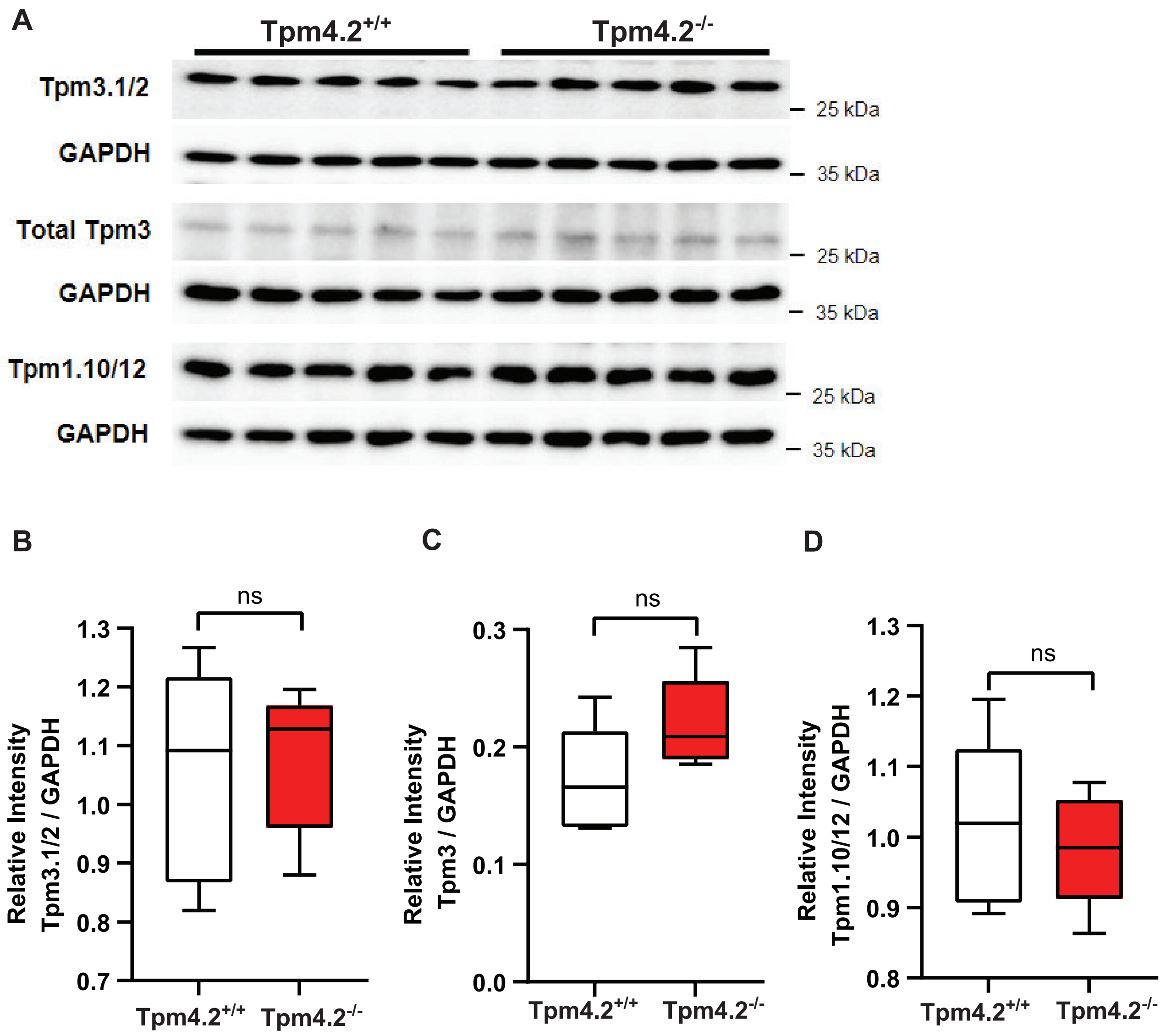

A
Tpm4.2+/+
Tpm4.2-/-
Tpm3.1/2
GAPDH
Total Tpm3
GAPDH
Tpm1.10/12
GAPDH
25 kDa
35 kDa
25 kDa
35 kDa
25 kDa
35 kDa
B
ns
Relative Intensity Tpm3.1/2 / GAPDH
1.3
1.2
1.1
1.0
0.9
0.8
0.7
Tpm4.2+/+
Tpm4.2-/-
C
ns
Relative Intensity Tpm3 / GAPDH
0.3
0.2
0.1
0.0
Tpm4.2+/+
Tpm4.2-/-
D
ns
Relative Intensity Tpm1.10/12 / GAPDH
1.3
1.2
1.1
1.0
0.9
0.8
Tpm4.2+/+
Tpm4.2-/-

Fig. S3

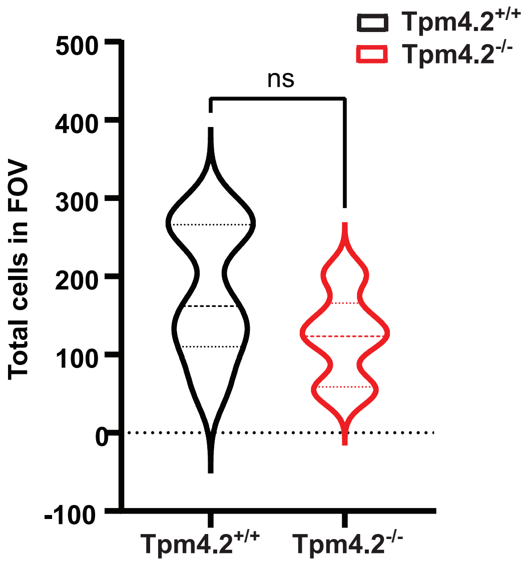

Tpm4.2+/+
Tpm4.2-/-
ns
500
400
300
200
100
0
-100
Total cells in FOV
Tpm4.2+/+
Tpm4.2-/-

Fig. S4

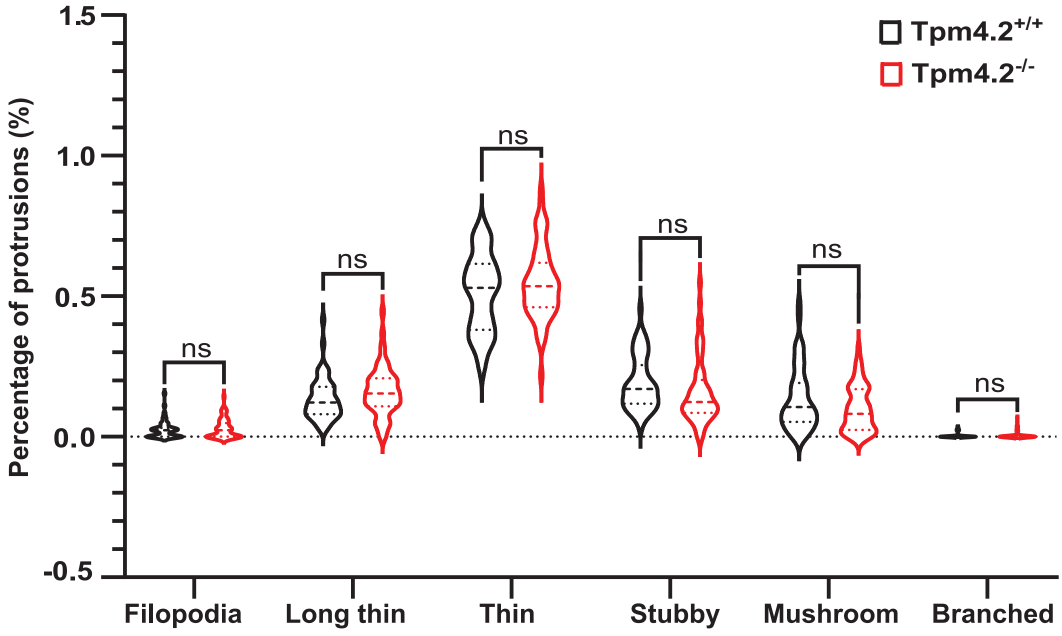

Tpm4.2+/+
Tpm4.2-/-
Percentage of protrusions (%)
1.5
1.0
0.5
0.0
-0.5
ns
ns
ns
ns
ns
ns
Filopodia
Long thin
Thin
Stubby
Mushroom
Branched

Fig. S5

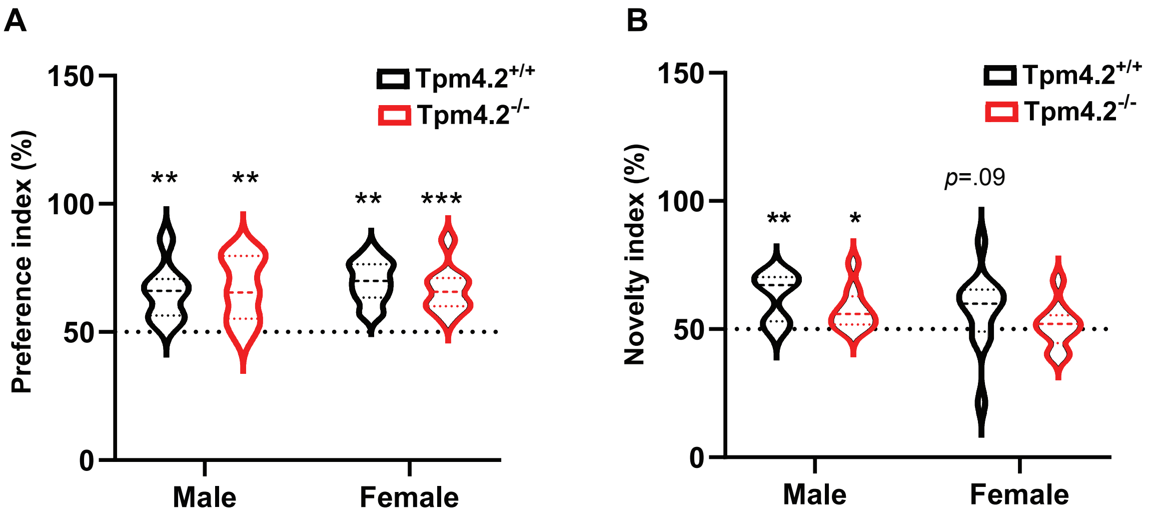
A
150
100
50
0
Preference index (%)
Tpm4.2+/+
Tpm4.2-/-
**
**
**
***
Male
Female
B
150
100
50
0
Novelty index (%)
Tpm4.2+/+
Tpm4.2-/-
**
*
p=.09
Male
Female

Fig. S6

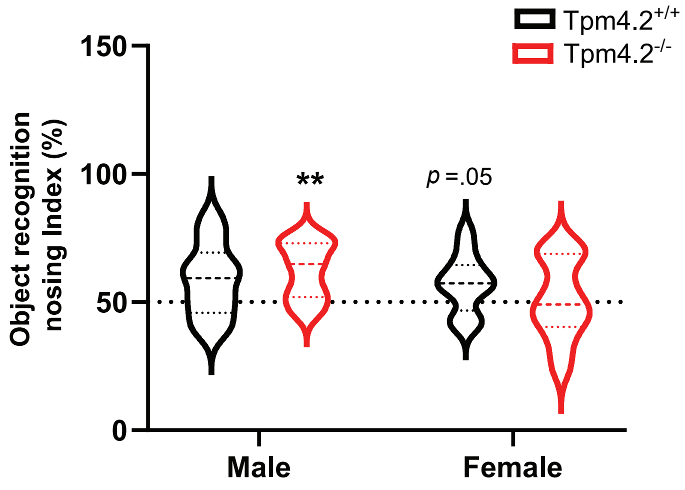
Tpm4.2+/+
Tpm4.2-/-
Object recognition nosing Index (%)
150
100
50
0
**
p =.05
Male
Female

Fig. S7

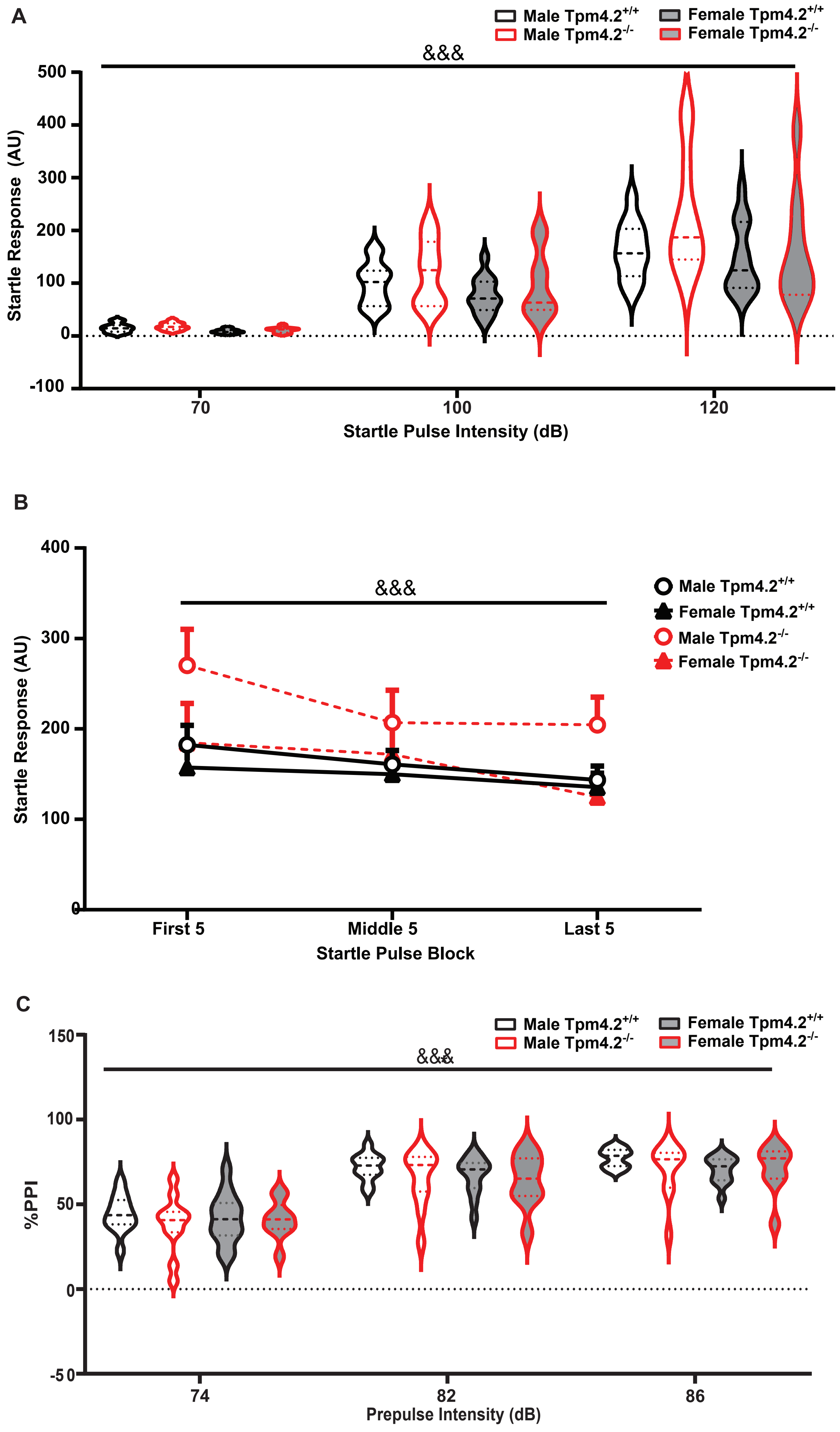
A
Male Tpm4.2+/+
Male Tpm4.2-/-
Female Tpm4.2+/+
Female Tpm4.2-/-
&&&
Startle Response (AU)
500
400
300
200
100
0
-100
70
100
120
Startle Pulse Intensity (dB)
B
400
300
200
100
0
&&&
Male Tpm4.2+/+
Female Tpm4.2+/+
Male Tpm4.2-/-
Female Tpm4.2-/-
Startle Response (AU)
First 5
Middle 5
Last 5
Startle Pulse Block
C
Male Tpm4.2+/+
Male Tpm4.2-/-
Female Tpm4.2+/+
Female Tpm4.2-/-
&&&
150
100
50
0
-50
%PPI
74
82
86
Prepulse Intensity (dB)

Table S1

**Tpm4.2$^{+/+}$**

| Olfactory Bulb | Hippocampus | Cerebellum | Cortex Anterior | Cortex Posterior |
|---|---|---|---|---|
| 1.659997439 | 0.698457785 | 0.658610751 | 0.255718227 | 0.310405001 |
| 0.338488807 | 0.330841497 | 0.494241986 | 0.335016314 | 0.317879974 |
| 0.521165498 | 0.43009832 | 0.808763004 | 0.325002436 | 0.298905311 |
| 0.783150183 | 1.017690336 | 0.892221198 | 0.584836146 | 0.463582373 |

**C57BL6**

| Olfactory Bulb | Hippocampus | Cerebellum | Cortex Anterior | Cortex Posterior |
|---|---|---|---|---|
| 1.264274197 | 0.673813753 | 1.268342784 | 1.006918731 | 0.72399428 |
| 0.987394417 | 0.925513045 | 1.376030633 | 0.806304645 | 0.830506028 |
| 0.65885429 | 0.264590199 | 1.197260274 | 0.31778112 | 0.501168289 |
| 0.847958562 | 0.454824515 | 1.347752063 | 0.902046309 | 0.885092615 |

Values displayed as relative signal intensity of Tpm4.2 / GAPDH detection.
